# Dual AAV gene therapy using laminin-linking proteins ameliorates muscle and nerve defects in LAMA2-related muscular dystrophy

**DOI:** 10.1101/2025.09.16.676550

**Authors:** Judith R. Reinhard, Shuo Lin, Eleonora Maino, Daniel J. Ham, Markus A. Rüegg

**Affiliations:** Biozentrum, University of Basel, 4056 Basel, Switzerland

## Abstract

Adeno-associated virus (AAV)-mediated gene replacement holds promise for treating genetic diseases but faces challenges due to AAV’s limited packaging capacity and potential immune responses to transgene products, especially in patients lacking endogenous protein. LAMA2-related muscular dystrophy (LAMA2 MD), a severe congenital disorder caused by loss of laminin-α2, presents both hurdles: the *LAMA2* gene exceeds AAV capacity, and severely affected patients do not produce the native protein. Here, we developed an AAV-based therapy using two engineered linker proteins derived from endogenously expressed components. These linker proteins restore laminin receptor binding and polymerization, enabling reassembly of a functional basement membrane. Dual AAV delivery of the linkers in a severe LAMA2 MD mouse model resulted in robust expression and significant improvements in muscle histology and function. Employing myotropic capsids enabled therapeutic efficacy at lower vector doses. However, muscle-specific targeting unmasked a LAMA2-related peripheral neuropathy. To address this, we expressed one linker under a muscle-specific promoter and the other under a ubiquitous promoter, delivered *via* AAV9 or AAV8. This approach achieved near-complete phenotypic restoration when administered neonatally and provided significant benefit when given at progressed disease stages. Our strategy offers a mutation-independent, size-compatible, and potentially immune-tolerable treatment for LAMA2 MD with broad clinical potential.

## INTRODUCTION

Laminins are a family of extracellular matrix (ECM) glycoproteins essential for the structural and functional integrity of tissues. As key components of basement membranes, they provide mechanical support, facilitate cell adhesion, and contribute to tissue organization and stability.^1^ Laminins are heterotrimeric proteins composed of one α, one β, and one γ chain, encoded by distinct genes. In mammals, five α (*LAMA1-5*), four β (*LAMB1-4*), and three γ (*LAMC1-3*) chains have been identified, which can assemble into at least 15 distinct isoforms in a tissue- and development-specific manner.^2^ The laminin-α2 chain, encoded by *LAMA2*, typically assembles with either a β1 or β2 chain and a γ1 chain, forming heterotrimers collectively referred to here as laminin-2X1 (α2, β1/2, γ1). The laminin-211 isoform, originally known as merosin, was identified to be predominantly expressed in Schwann cells and striated muscle.^3^

Recessive loss-of-function mutations in *LAMA2*, which encodes the laminin-α2 chain, cause LAMA2-related muscular dystrophy (LAMA2 MD), the disease was first identified as congenital early-onset severe muscular dystrophy.^4^ Today, it is clear that the disease exists in two forms. The severe, early-onset congenital form presents within the first six months of life and is typically caused by biallelic null mutations resulting in complete absence of laminin-α2. The milder, late-onset form is often associated with hypomorphic mutations that allow some residual protein expression.^5,6^ Clinically, LAMA2 MD manifests with progressive muscle wasting, motor impairments, respiratory insufficiency, and, in severe cases, early mortality.^7–9^ Most early-onset patients do not achieve independent ambulation, while late-onset patients may attain and later lose the ability to walk.^6,7,10^ Despite the clinical burden, no disease-modifying treatments are currently available.

Several mouse models recapitulate the human disease with varying severity. The *dy^2J^*/*dy^2J^* model mimics a late-onset phenotype and maintains normal survival, whereas *dy^W^*/*dy^W^*and *dy^3K^*/*dy^3K^* mice exhibit severe early-onset disease with severe muscle wasting and strongly reduced lifespan.^11^ In addition to muscular dystrophy, all mouse models consistently present with pronounced peripheral neuropathy.^12^ In human patients, peripheral neuropathy is less apparent. However, some studies have reported abnormalities in peripheral nerve conduction and morphology, supporting peripheral nerve involvement in the disease.^12^

Gene therapy offers a promising avenue for treating LAMA2 MD. However, the large size of the LAMA2 coding sequence exceeds the packaging capacity of adeno-associated virus (AAV) vectors, precluding direct gene replacement. Furthermore, immune responses to transgene products remain a significant challenge in neuromuscular gene therapy, especially in immunologically primed or cross-reactive immunologic material (CRIM)-negative patients.^13–16^ In LAMA2 MD, severe early-onset cases present with extensive inflammatory infiltration,^17–19^ which may exacerbate immune responses to exogenous proteins - particularly bacterial components, such as Cas9 or split-intein domains used in gene editing or dual-vector strategies. Additionally, the intracellular assembly of laminin heterotrimers via coiled-coil domains complicates the design of functional mini-laminins or fusion constructs, as alterations at junction sites may interfere with proper trimerization and ECM incorporation.

To overcome these barriers, we developed a mutation-independent gene therapy strategy that leverages the compensatory upregulation of laminin-α4 observed in LAMA2 MD patients.^20,21^ In the absence of laminin-α2, muscle tissue express laminin-4X1 (α4,β1/2,γ1), which is properly assembled and secreted, but functionally inadequate. Laminin-4X1 cannot polymerize or bind α-dystroglycan and α7β1-integrin that normally anchor laminin-2X1 to the muscle fiber surface.^21^ To restore basement membrane function, we engineered two linker proteins composed of endogenously expressed domains of other ECM components. These linkers enable receptor binding and polymerization of laminin-4X1, effectively reconstituting a functional basement membrane when co-delivered via dual AAV vectors.

In the severe *dy^W^*/*dy^W^* mouse model, expression of the linkers via systemic AAV9 or AAV8 application led to widespread incorporation into the basement membrane and robust improvements in muscle histology, function, and transcriptional profiles. Importantly, due to their secreted nature, the linkers achieve broad tissue distribution without requiring increased total AAV doses. Using myotropic AAV capsids enabled efficient transduction at lower doses.

In addition to the muscular dystrophy, we also targeted the peripheral neuropathy by expressing one linker by a ubiquitous promoter. This dual-tissue targeting ameliorated both muscular dystrophy and peripheral neuropathy. Importantly, the therapy was effective when administered neonatally and retained therapeutic benefit when delivered at later disease stages. Thus, our approach bypasses the limitations imposed by vector size and immune risk, offering a potentially translatable, mutation-independent treatment for LAMA2 MD.

## RESULTS

### AAV-mediated muscle-specific expression of linker proteins improves muscle histology, body weight and muscle function in *dy^W^*/*dy^W^* mice

Since the coding sequence of laminin-α2 is too large for gene replacement via AAVs, we aimed to restore laminin-α2 function by leveraging laminin-4X1, which is present in the basement membrane of laminin-α2-deficient patients and mouse models.^20,21^ To achieve this, we used mini-agrin (mag) to anchor laminin-4X1 to the sarcolemma via α-dystroglycan, and αLNNd to enable laminin-4X1 polymerization (Figure 1A). Mag is a miniaturized form of the ECM proteoglycan agrin. It includes the N-terminal agrin (NtA) domain, which binds to the coiled-coil region of laminins,^22^ and a C-terminal part that binds with high affinity to α-dystroglycan.^23^ The C-terminal domains lack the amino acid inserts A/y and B/z, which are required for acetylcholine receptor clustering.^24,25^ The second linker, αLNNd, consists of the LN domain of laminin-α1 fused to the laminin-γ1-binding domain of nidogen-1.^26^ Both engineered linkers contained their endogenous signal peptides to ensure proper secretion (Figure 1B).

**Figure 1.**
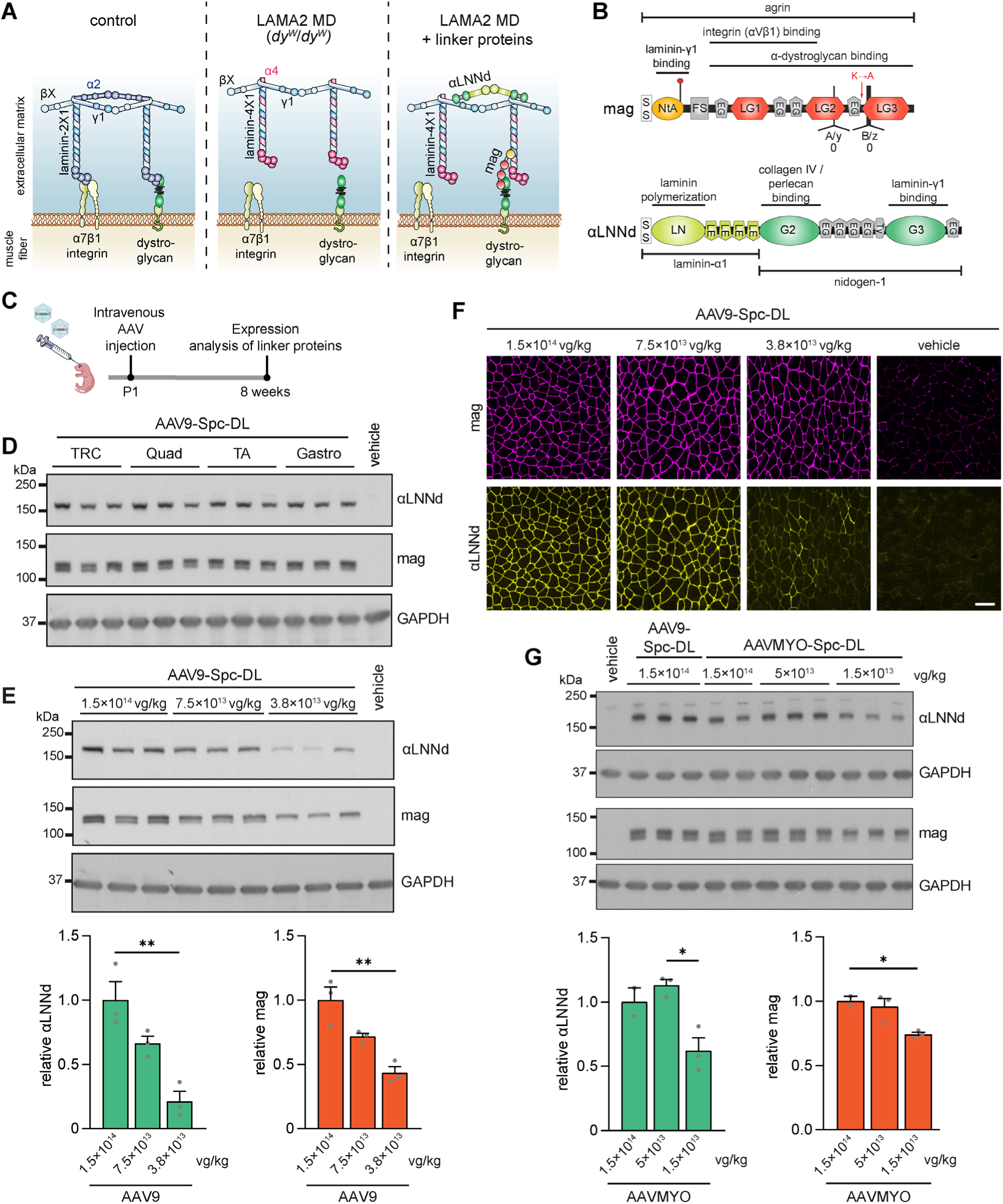
AAV-mediated expression of linker proteins to restore laminin receptor anchorage and polymerization. (**A**) Schematic representation of laminin heterotrimers containing laminin-α2 [laminin-2X1 (α2, β1/2, γ1)] or laminin-α4 [laminin-4X1 (α4, β1/2, γ1)], and the linker proteins mag and αLNNd in the muscle basement membrane. (**B**) Domain structure of the linker proteins: mag is a fusion protein comprising the N-terminal and C-terminal regions of agrin, that includes the domains mediating binding to laminins and α-dystroglycan. αLNNd is a fusion protein combining the LN domain and LE modules of laminin-α1 (conferring polymerization activity) with the C-terminal domains of nidogen-1 (providing laminin-γ1 and collagen IV binding sites). (**C**) Experimental design for panels D-G: C57BL/6 mice were injected intravenously at postnatal day 1 (P1) with the indicated combination of AAV vectors. Muscle tissue was analyzed at 8 weeks of age. (**D-E**) Western blot analysis to detect αLNNd and mag in lysates from various muscles (**D**) or triceps muscle (**E**) of mice injected with AAV9-Spc-αLNNd and AAV9-Spc-mag (referred to as AAV9-Spc-DL). Each mouse received 1.5×10¹⁴ vg/kg per construct (**D**) or the indicated dose (**E**). GAPDH was used as loading control. (**F**) Immunofluorescence images of triceps cross-sections stained for αLNNd and mag from mice injected with AAV9-Spc-DL at the indicated doses. (**G**) Western blot analysis to detect αLNNd and mag in lysates from triceps of mice injected with AAV9-Spc-DL or AAVMYO-Spc-DL at the indicated doses. Relative amounts of αLNNd and mag were normalized to the expression of their respective linker at the highest AAV9 dose. Data are presented as mean ± SEM. \**P* < 0.05 and \*\**P* < 0.01 by one-way ANOVA with Bonferroni post hoc test. Scale bar: 100 μm. *n* = 2-3 mice per group.

To assess the feasibility of AAV-mediated delivery, we generated AAV9 vectors encoding codon-optimized versions of human mag (mag) and human αLNNd (αLNNd). Given the large size of αLNNd (4.1 kb), we used the synthetic, small muscle-specific promoter Spc5.12 (Spc)^27,28^ for both constructs. To enable detection of mag, we generated a polyclonal rabbit antiserum by immunizing rabbits with His-tagged mag, which was expressed in HEK293 cells and subsequently purified from conditioned medium (Figure S1A). The resulting serum recognized mag with high specificity and sensitivity (Figure S1B). αLNNd was detected using commercial antibodies, raised against the N-terminal domain of human laminin-α1. As laminin-α1 is not expressed in mouse skeletal muscle,^29^ laminin-α1 antibodies were used for immunohistochemistry and Western blot analysis. To test for linker expression, we injected a mixture of AAV9-Spc-αLNNd and AAV9-Spc-mag, hereafter referred to as AAV9-Spc-DL (DL for “double linker”), into the temporal vein of wild-type mice at postnatal day 1 (P1) and determined the protein amount in tissues at 8 weeks (experimental scheme in Figure 1C). Each mouse received a dose of 1.5×10¹⁴ vector genomes/kilogram (vg/kg) per construct. Analysis of multiple muscles and the liver showed high transduction efficiency, as indicated by the quantification of vg per mouse genome (Figure S1C), and of the amount of the linker proteins in various muscles (Figure 1D). To evaluate dose-dependent expression, we lowered the AAV dose up to 4-fold (Figure 1E, F). Reducing the dose by half only slightly decreased the linker protein amount and basement membrane localization remained homogenous, suggesting that injection of 7.5×10^13^ vg/kg is still sufficient to saturate muscles with linker proteins. Additional halving of the dose led to a clear decrease of the signal in Western blots (Figure 1E) and to non-homogenous appearance in immunohistochemistry, particularly for αLNNd (Figure 1F).

Next, we assessed whether the myotropic AAV variant AAVMYO^30^ would increase muscle delivery of the linker proteins. At a dose of 5×10^13^ vg/kg of AAVMYO, the amount of mag and αLNNd was not significantly different to what was achieved with AAV9 at 1.5×10^14^ vg/kg (Figure 1G). At a dose of 1.5×10^13^ vg/kg, AAVMYO-delivered linker proteins were still at 74% (mag) and 62% (αLNNd) of the levels observed at maximal dose (Figure 1G).

Interestingly, the maximal amounts detected with AAVMYO were not higher than those achieved with 1.5×10^14^ vg/kg AAV9 (Figure 1G, top), suggesting that we reach saturation at the highest doses irrespective of the delivery AAV capsid.

Since the therapeutic effects of mouse-derived linker proteins have previously been evaluated in LAMA2 MD mouse models through transgenic expression,^21,31^ we aimed to assess the efficacy of AAV-mediated delivery of the corresponding mouse linker protein sequences. To this end, we injected *dy^W^*/*dy^W^* mice at P1 with a high, saturating dose of 1.5×10^14^ vg/kg of AAV9 carrying the coding sequence for the mouse versions of the linker proteins (AAV9-Spc-m.DL). Treatment effects were assessed until 8 weeks of age (Figure 2A). At this age *dy^W^*/*dy^W^* mice present with a very severe disease phenotype, but approximately 60% of the untreated mice still survive.^21^ This way, we wanted to ensure that we did not particularly select for long-survivors in the non-treated cohort. Histological analysis (H&E stainings) of muscle cross-sections from untreated *dy^W^*/*dy^W^* mice revealed characteristic features of severe muscular dystrophy, including pronounced fiber size variation, extensive fibrosis, mononuclear cell infiltration, and presence of centralized myonuclei in both triceps (Figure 2B) and diaphragm muscle (Figure S2A). In contrast, muscle histology was markedly improved in AAV9-Spc-m.DL-treated *dy^W^*/*dy^W^* mice. Notably, the median triceps muscle fiber diameter was restored to control levels (Figure 2C) and diaphragm thickness was significantly higher than in untreated mice (Figure S2A). In line with the histological improvements, treated mice displayed visibly enhanced body size and overall appearance (Video S1). These findings demonstrate that AAV-mediated expression of mouse linker proteins achieves therapeutic efficacy comparable to that of transgenic expression.

**Figure 2.**
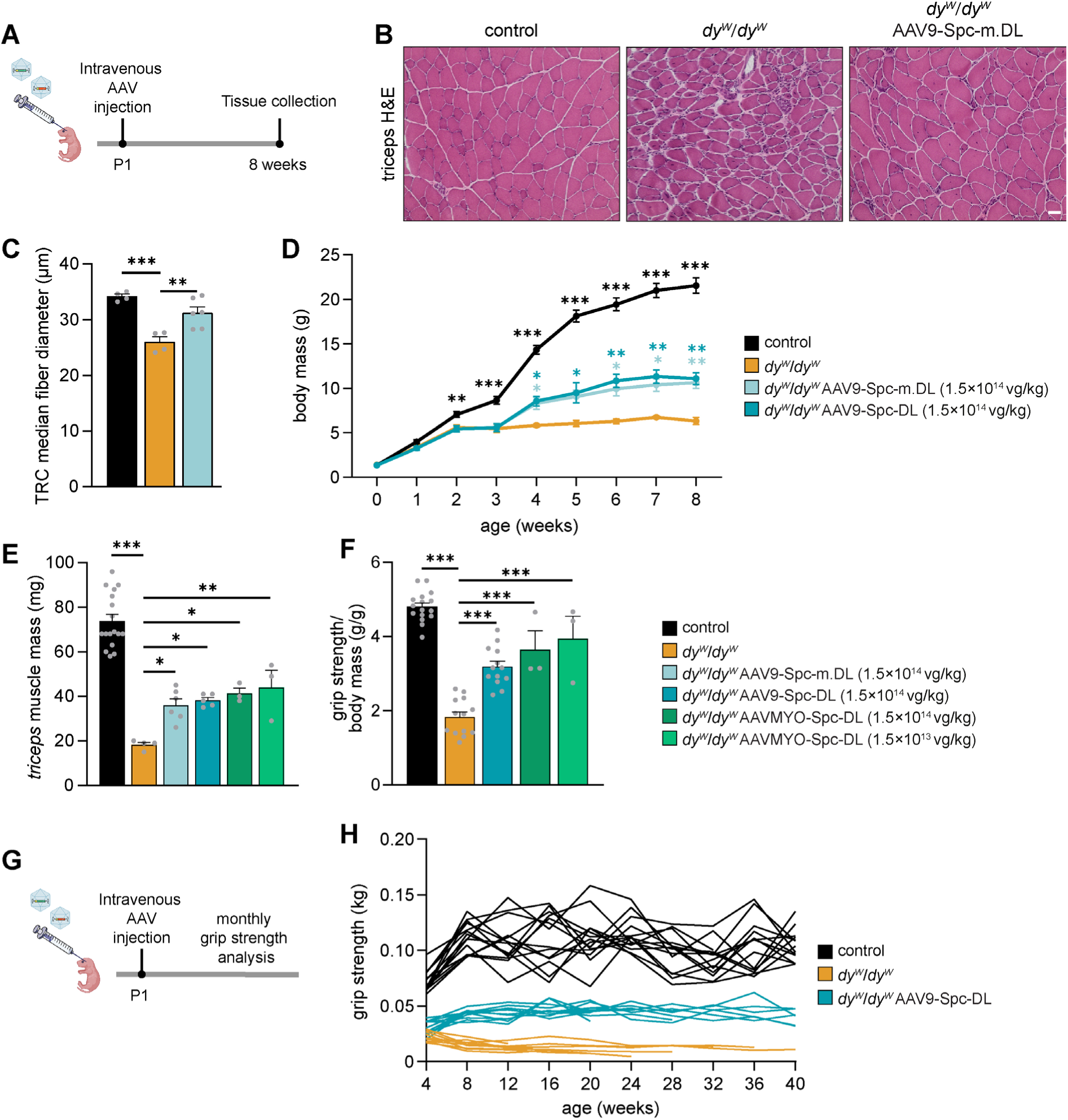
AAV-mediated, muscle targeted expression of linker proteins for amelioration of muscular dystrophy in *dy^W^*/*dy^W^* mice. (**A**) Experimental design for panels B-D: *dy^W^*/*dy^W^* mice were injected intravenously at P1 with the indicated combination of AAV vectors. Muscle tissue was analyzed at 8 weeks of age. (**B-C**) Histological assessment of triceps muscles from non-injected healthy control, non-treated *dy^W^*/*dy^W^* mice, or *dy^W^*/*dy^W^* mice treated with AAV9-Spc-m.αLNNd and AAV9-Spc-m.mag (referred to as AAV9-Spc-m.DL) at a dose of 1.5×10¹⁴ vg/kg per construct. (**B**) Representative pictures of H&E-stained triceps cross-sections. (**C**) Quantification of median fiber diameters in triceps cross-sections. (**D**) Body mass curves of non-injected healthy control and *dy^W^*/*dy^W^* mice, as well as *dy^W^*/*dy^W^* mice treated with AAV9-Spc-m.DL or the corresponding human linker constructs (AAV9-Spc-DL). (**E**) Triceps muscle mass at 8 weeks of age of the different groups. (**F**) Forelimb grip strength normalized to body mass of the different groups. (**G**) Experimental design for panel H: *dy^W^*/*dy^W^* mice were injected intravenously at P1 with AAV9-Spc-DL (1.5×10¹⁴ vg/kg per construct) and evaluated monthly for grip strength. (**H**) Longitudinal assessment of individual forelimb grip strength. Discontinuous lines indicate that the mouse died during the course of the study. Data are presented as mean ± SEM. \**P* < 0.05, \*\**P* < 0.01, \*\*\**P* < 0.001 by one-way ANOVA with Dunnett’s post hoc test, comparing each group to untreated *dy^W^*/*dy^W^* mice. Scale bar: 50 μm. *n* = 3-17 mice per group.

The amino acid sequence of the linker proteins is highly conserved between mouse and human, reaching 84% similarity for mag and 87% similarity for αLNNd. To determine whether the human linker proteins are functionally equivalent to their mouse homologs, we conducted a direct comparison. Mice injected at P1 with AAV9 vectors encoding either the mouse (m.DL) or the human (DL) versions of the linker proteins showed comparable weight gain starting three weeks post-injection (Figure 2D), ultimately reaching approximately a two-fold increase in body weight by 8 weeks of age compared to untreated *dy^W^*/*dy^W^* mice. Similarly, the mass of various skeletal muscles was comparably increased in both treated groups (Figure S2B). These results provide strong evidence that the human versions are as effective as their mouse homologs. Given the necessity of using human sequences for future clinical translation, all subsequent experiments were conducted using the human linker proteins.

To determine whether the efficiency achieved with the maximal dose of AAV9 (1.5×10¹⁴ vg/kg) could be further enhanced, we next evaluated the use of AAVMYO for linker protein delivery. To assess both its potential superiority and potency, we injected *dy^W^*/*dy^W^* mice with AAVMYO-Spc-DL at the maximal dose (1.5×10^14^ vg/kg) or a ten-fold lower dose. Consistent with protein expression levels detected by Western blot analysis across the dose range, both triceps muscle mass and forelimb grip strength at 8 weeks of age showed significant improvement in treated *dy^W^*/*dy^W^*mice. These improvements were comparable to those observed with high-dose of AAV9 and both high- and low-dose AAVMYO (Figure 2E, F). These results indicate that the use of AAVMYO would allow a ten-fold dose reduction without loss of efficacy, and support the notion that both, linker protein amount and therapeutic effect approach saturation at the doses tested.

To assess the long-term durability of the treatment, we performed monthly forelimb grip strength measurements (Figure 2G). Treated *dy^W^*/*dy^W^*mice exhibited increased strength beginning at 4 weeks of age, and the grip strength remained stable up to the age of 40 weeks (Figure 2H). Note that only one untreated *dy^W^*/*dy^W^* mouse survived up to 40 weeks, while 50% of the treated mice survived past 40 weeks (Figure 2H). The sustained therapeutic effect of AAV-mediated linker protein expression was further supported by analysis of the triceps muscle from a treated *dy^W^*/*dy^W^* mouse that survived for two years. Even at this advanced age, histological improvements were evident, including increased myofiber size and a reduced infiltration of mononuclear cells (Figure S2C). Moreover, the basement membrane surrounding all muscle fibers remained positive for mag (Figure S2D). Similar to the pattern observed at the low dose (Figure 1F), αLNNd expression appeared more mosaic, potentially due to the larger size of the construct or the lower affinity of the detection antibody (Figure S2D).

### αLNNd, but not mini-agrin, ameliorates peripheral neuropathy in *dy^W^*/*dy^W^* mice

While body weight, muscle histology and function were significantly improved, the use of muscle-specific promoters to drive linker protein expression restricted the therapeutic effect to muscle and unmasked a progressive peripheral neuropathy that ultimately led to complete hindlimb paralysis (Video S2). In all laminin-α2-deficient mouse models, peripheral neuropathy predominantly affects the hindlimbs and is characterized by defects in axonal sorting and myelination.^12^ Notably, skeletal muscle-specific expression of *Lama2* also fails to correct the peripheral neuropathy in *dy^W^*/*dy^W^*mice ^32^. In contrast, ubiquitous transgenic expression of both linker proteins prevents hindlimb paralysis in *dy^W^*/*dy^W^*and *dy^3K^*/*dy^3K^* mice.^31^ Hence, the linker proteins are also effective in improving peripheral nerve pathology when expressed ubiquitously.

To determine whether both linker proteins are required to prevent hindlimb paralysis, we analyzed *dy^W^*/*dy^W^*mice transgenically expressing only one linker protein under the control of the ubiquitous CAG promoter. Strikingly, *dy^W^*/*dy^W^*mice expressing solely the αLNNd linker protein (*dy^W^*/*dy^W^* CAG-αLNNd) did not develop hindlimb paralysis (Video S3), whereas those expressing only mag (*dy^W^*/*dy^W^*CAG-mag) showed no improvement (Video S4). Locomotor function analysis of 8-week-old mice confirmed that *dy^W^*/*dy^W^* CAG-αLNNd mice, but not *dy^W^*/*dy^W^* CAG-mag, achieved a near-normal gait speed compared to *dy^W^*/*dy^W^* mice (Figure 3A). Consistent with these functional improvements, sciatic nerve histology revealed normal axonal sorting and myelination in *dy^W^*/*dy^W^* CAG-αLNNd mice, whereas *dy^W^*/*dy^W^* mice showed large bundles or islands of “naked” axons containing many large caliber axons (Figure S3A).

**Figure 3.**
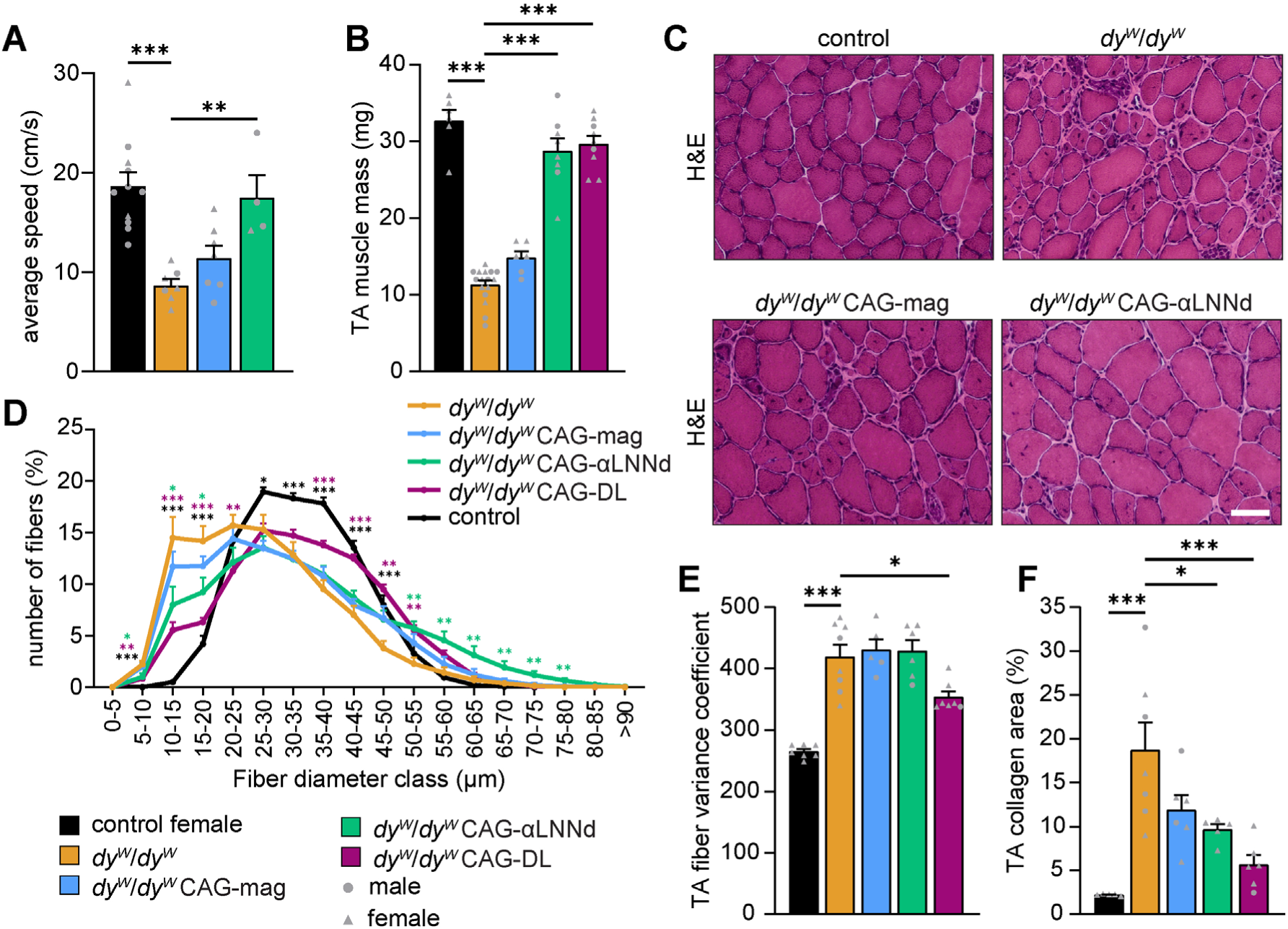
Effect of ubiquitous transgenic expression of αLNNd and mag on hindlimb neuropathy and muscular dystrophy in *dy^W^*/*dy^W^* mice. (**A**) Quantitative analysis of locomotor speed in 8-week-old mice using a gait analysis system. Groups include healthy controls, *dy^W^*/*dy^W^*mice, and *dy^W^*/*dy^W^* transgenic mice ubiquitously expressing mag or αLNNd under the control of the CAG promoter (CAG-mag or CAG-αLNNd). (**B**) Tibialis anterior (TA) muscle mass at 8 weeks of age in control mice, *dy^W^*/*dy^W^* mice, and *dy^W^*/*dy^W^* transgenic mice expressing individual linker proteins or both in combination (CAG-DL). (**C-F**) Histological analysis of TA muscles from 8-week-old controls, *dy^W^*/*dy^W^* mice, and *dy^W^*/*dy^W^* transgenic mice as indicated. (**C**) Representative images of H&E-stained TA cross-sections. (**D**) Distribution of muscle fiber diameters in TA cross-sections. (**E**) Variance coefficient of TA fiber diameters. (**F**) Quantification of muscle fibrosis based on Sirius Red-positive area in TA cross-sections. Data are presented as mean ± SEM. \**P* < 0.05, \*\**P* < 0.01, \*\*\**P* < 0.001 by one-way ANOVA with Dunnett’s post hoc test, comparing each group to *dy^W^*/*dy^W^*mice. Scale bar: 100 μm. *n* = 4-11 mice per group.

In prior experiments, we observed that muscle-specific expression of the two linker proteins could not fully restore hindlimb muscle weight in *dy^W^*/*dy^W^*mice ^21^, likely due to secondary muscle atrophy caused by the peripheral neuropathy. To address this, we analyzed tibialis anterior (TA) muscles from *dy^W^*/*dy^W^*CAG-αLNNd and *dy^W^*/*dy^W^* CAG-mag mice. While muscle mass was restored in *dy^W^*/*dy^W^* CAG-DL (double transgenic for both linkers; DL for “double linker”) and *dy^W^*/*dy^W^*CAG-αLNNd mice, *dy^W^*/*dy^W^* CAG-mag mice retained the characteristic atrophy-linked muscle mass reduction (Figure 3B). Partial histological improvements could be observed in TA muscle with either linker protein alone (Figure 3C), consistent with prior muscle-specific transgenic studies ^21^. Similarly, expression of both linker proteins was required to restore fiber size distribution comparable to control female mice (Figure 3D), while αLNNd expression alone reduced the proportion of small fibers but increased the number of exceptionally large fibers. Importantly, significant reduction in fiber size variability also required the presence of both linker proteins, as neither linker alone improved this parameter (Figure 3E). Assessment of fibrosis, determined by the collagen area in Sirius Red staining, showed a decrease with either linker protein, with the greatest reduction observed again when both linkers were combined (Figure 3F).

Collectively, and in agreement with previous findings, these results demonstrate the additive benefit of the two linker proteins on muscle histology,^21^ whereas αLNNd alone is sufficient to ameliorate the peripheral neuropathy.

To explore the mechanistic basis for this differential effect, we assessed laminin-α4 expression in peripheral nerves. In wild-type mice, laminin-α4 localizes primarily to the perineurium, whereas laminin-α2 is the predominant laminin α-chain in the endoneurium.^31,33^ In *dy^W^*/*dy^W^* mice, laminin-α4 is also detected in the endoneurium (Figure S3B).^31^ Strikingly, laminin-α4 staining was markedly increased in the endoneurium of *dy^W^*/*dy^W^*CAG-αLNNd mice but remained unchanged in *dy^W^*/*dy^W^*CAG-mag mice (Figure S3B). Western blot analysis corroborated this finding, confirming elevated laminin-α4 levels in *dy^W^*/*dy^W^* CAG-αLNNd mice (Figure S3C). These findings demonstrate that αLNNd sequesters laminin-4X1 within the endoneurial basement membrane. Given that Schwann cells - in contrast to muscle fibers - express the laminin-4X1 receptor integrin-α6β1, ^34,35^ the linkage of laminin-4X1 to α-dystroglycan via mag may not be essential in peripheral nerves. Instead, the ability of αLNNd to polymerize laminin-4X1 may enable a proper basement membrane formation and allow its engagement with integrin-α6β1, providing a mechanistic explanation for the functional restoration observed in *dy^W^*/*dy^W^* CAG-αLNNd mice.

### Shortening of αLNNd for efficient AAV9-mediated expression in muscle and nerve

To enable AAV-mediated ubiquitous expression of αLNNd, we engineered a truncated variant, αLNNdΔG2, by removing the nidogen-G2 domain and two adjacent EGF-like domains (Figure 4A). This truncation preserves laminin polymerization activity^36^ while reducing the coding sequence to comply with AAV packaging limits when using a larger promoter. To dissect the influence of the promoter and the construct size to expression levels, we generated AAV constructs encoding full-length αLNNd or αLNNdΔG2, driven by either the muscle-specific Spc5.12 promoter or the ubiquitous CBh promoter.^37^. Protein expression was evaluated four weeks post-injection (Figure 4B). Western blot analysis of TA muscle lysates showed higher αLNNdΔG2 levels compared to αLNNd when driven by the Spc5.12 promoter, with expression levels further increased by the CBh promoter (Figure 4C, D). To account for potential size-dependent transfer efficiency differences in Western blot analysis, we performed LC-MS using peptides shared by both constructs. LC-MS confirmed a similar expression pattern, albeit with attenuated absolute fold changes compared to Western blot data (Figure 4E). The increase in protein amount also correlated with elevated transcript levels (Figure 4F). Most importantly, CBh-driven αLNNdΔG2 - unlike the Spc5.12-driven construct – localized to the basement membrane of the sciatic nerve, including both the peri- and endoneurium (Figure 4G).

**Figure 4.**
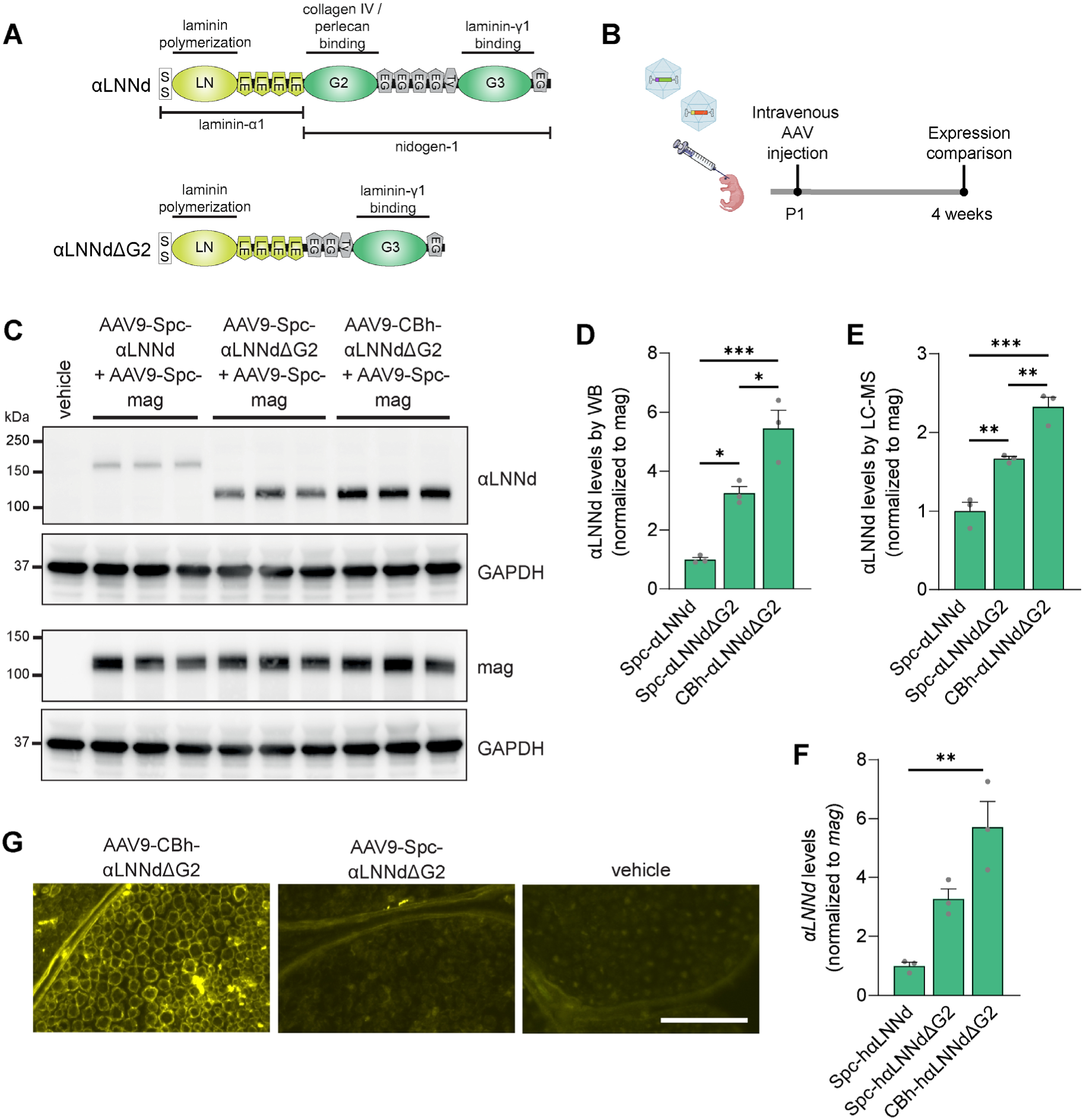
Shortening of αLNNd to allow CBh promoter-driven expression in peripheral nerve. (**A**) Schematic representation of the protein domains present in full-length αLNNd and its shortened version αLNNdΔG2. (**B**) Experimental design for panels C-F: C57BL/6 mice were injected intravenously at P1 with PBS (vehicle) or with the indicated combination of AAV vectors (1.5×10¹⁴ vg/kg per construct). Tissue was analyzed at 4 weeks of age. (**C-E**) Analysis of αLNNd, αLNNdΔG2, and mag protein amounts in triceps muscle lysates by Western blot (**C-D**) or LC-MS (**E**), and transcript levels by qPCR (**F**). (**C**) Western blot detection of mag, αLNNd, or αLNNdΔG2. Antibodies against the LN domain of laminin-α1 were used to detect both αLNNd variants. GAPDH served as a loading control. (**D**) Quantification of protein levels from Western blot analysis. (**E**) Quantification of protein abundance by LC-MS. (**F**) Quantification of transcript levels by qPCR. (**G**) Immunofluorescence images of sciatic nerve cross-sections stained for αLNNd or αLNNdΔG2 using antibodies against the LN domain of laminin-α1. Data are presented as mean ± SEM. \**P* < 0.05, \*\**P* < 0.01, \*\*\**P* < 0.001 by one-way ANOVA with Bonferroni’s post hoc test. *n* = 3 mice per group.

Given that the myotropic AAVMYO achieves higher expression of the linker proteins in skeletal muscle than AAV9 at several-fold lower doses (Figure 1G), we evaluated its potential to target peripheral nerves. At the dose of 1.5×10^13^ vg/kg, αLNNdΔG2 could not be detected in the endoneurium of the peripheral nerve (Figure S4A). We next tested AAVMYO2, another myotropic capsid generated through shuffling and peptide display, that additionally de-targets the liver ^38^. Even at a tenfold higher dose (1.5×10^14^ vg/kg), AAVMYO2-delivered αLNNdΔG2 protein could not be detected in the peripheral nerve (Figure S4B). As expected, AAVMYO2 clearly outperformed AAV9 in skeletal muscle at lower doses (Fig. S4C-E).

In summary, αLNNdΔG2 delivered by AAV9 at a dose of 1.5×10^14^ vg/kg using the CBh promoter localizes to both the muscle and peripheral nerve basement membrane, suggesting that it may ameliorate pathology in both tissues following AAV administration. Furthermore, this construct showed markedly improved expression in muscle compared to Spc-αLNNd, potentially improving durability and enabling therapeutic efficacy at lower doses. We also concluded that AAVMYO and AAVMYO2 are not suitable vectors to deliver αLNNdΔG2 to the peripheral nerve and therefore excluded them from further evaluation.

### Early treatment of *dy^W^*/*dy^W^* mice with ubiquitously expressed αLNNdΔG2 and muscle-specific mag reduces all disease hallmarks

Having established that αLNNdΔG2 is efficiently expressed in peripheral nerves using AAV9 and prevents the peripheral neuropathy when expressed transgenically, we next tested its therapeutic efficiency when delivered to *dy^W^*/*dy^W^* mice via AAV9. To this end, we combined AAV9 vectors encoding mag under the control of the Spc5.12 promoter (Spc-mag; Figure 5A) with an equal amount of AAV9 encoding αLNNdΔG2 driven by the CBh promoter (CBh-αLNNdΔG2; Figure 5A). The AAV9 mixture (referred to as AAV9-Spc/CBh-DL) was then intravenously administered to *dy^W^*/*dy^W^* mice at P1 at a dose of 1×10^14^ vg/kg per construct, and tissues were analyzed at 8 weeks of age (Figure 5B). Histological analysis of TA muscle revealed a strong reduction in the dystrophic hallmarks (Figure 5C, Figure S5A), including decreased fibrosis as demonstrated by Sirius Red staining (Figure 5C). This improvement in fibrosis was further confirmed by highly significant reduction in collagen content, measured by hydroxyproline quantification (Figure 5D). Muscle fiber size distribution shifted markedly towards larger fibers (Figure 5E), accompanied by increases in median fiber diameter (Figure 5F) and fiber number (Figure 5G). Despite these improvements, a high proportion of fibers retained centralized nuclei (Figure 5H). TA muscle of treated *dy^W^*/*dy^W^* mice also showed little to no signs of inflammation as determined by staining using antibodies against the macrophage marker F4/80 (Figure 5I). Quantitative assessment of the disease amelioration by the linker proteins was very similar in triceps muscle (Figure S5B–E).

**Figure 5.**
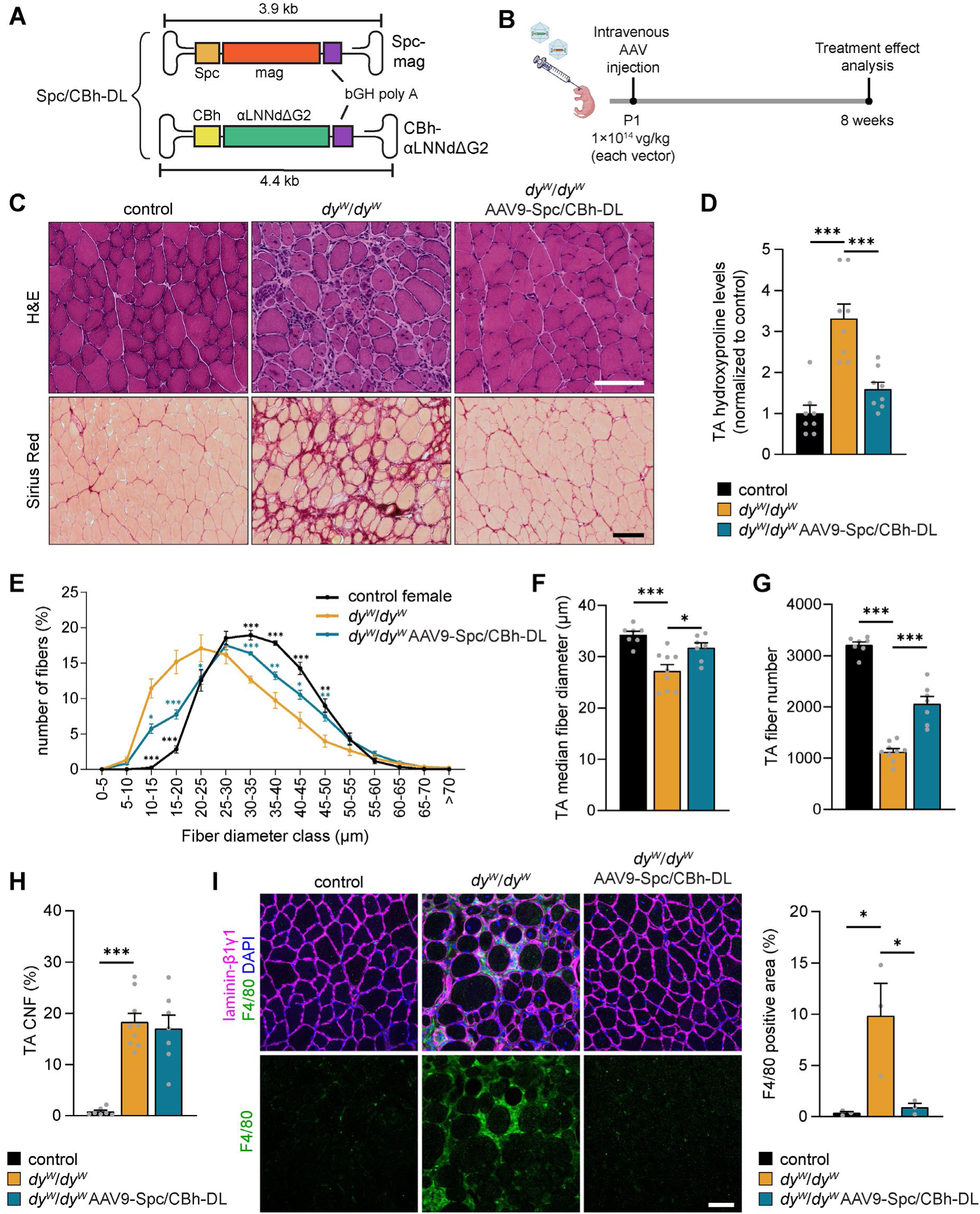
Hindlimb muscle histology upon AAV-mediated ubiquitous expression of αLNNdΔG2 combined with muscle-specific expression of mag in *dy^W^*/*dy^W^*mice. (**A**) Schematic illustration of vector constructs used in the Spc/CBh-DL treatment. Expression of mag is driven by the muscle-specific Spc promoter, while αLNNdΔG2 is expressed under the control of the ubiquitous CBh promoter. (**B**) Experimental design for data shown in Figures 5-7. *dy^W^*/*dy^W^* mice were injected intravenously at P1 with a mixture of AAV9 vectors: one packaged with Spc-mag the other packaged with CBh-αLNNdΔG2. This combination is referred to as AAV9-Spc/CBh-DL. Each construct was administered at a dose of 1×10¹⁴ vg/kg. Treatment effects were analyzed at 8 weeks of age. (**C**) Representative images of H&E- and Sirius Red-stained TA muscle cross-sections from non-injected healthy controls, untreated *dy^W^*/*dy^W^* mice, and *dy^W^*/*dy^W^* mice treated with AAV9-Spc/CBh-DL. (**D**) Quantification of muscle fibrosis based on hydroxyproline content measured by amino acid analysis in TA lysates. (**E-H**) Quantitative analysis of TA muscle histology in non-injected healthy controls, untreated *dy^W^*/*dy^W^* mice, and *dy^W^*/*dy^W^* mice treated with AAV9-Spc/CBh-DL. (**E**) Distribution of muscle fiber diameters. (**F**) Median muscle fiber diameters. (**G**) Total muscle fiber number. (**H**) Percentage of fibers with centralized nuclei. (**I**) Immunofluorescence images of TA cross-sections stained for laminins (β1γ1-chains) and F4/80 (macrophage marker), along with quantification of F4/80-positive area. Data are presented as mean ± SEM. \**P* < 0.05, \*\**P* < 0.01, \*\*\**P* < 0.001 by one-way ANOVA with Dunnett’s post hoc test, comparing each group to untreated *dy^W^*/*dy^W^*mice. Scale bar: 100 μm. *n* = 3-9 mice per group.

To assess the global transcriptional effects of the treatment, we performed bulk RNA-seq on TA muscle from 8-week-old mice. We included six wild-type controls, six untreated *dy^W^*/*dy^W^* mice and six *dy^W^*/*dy^W^*mice treated with AAV9-Spc/CBh-DL mice (three males and three females per group). In addition, we included three healthy control mice injected with AAV9-Spc/CBh-DL to evaluate potential off-target effects of the treatment. Unbiased sample correlation analysis revealed a clear separation between untreated *dy^W^*/*dy^W^* mice and treated *dy^W^*/*dy^W^*mice, while wild-type and AAV9-Spc/CBh-DL-injected wild-type mice were intermixed (Figure S6A). This indicates a strong transcriptional effect of the linker protein treatment in *dy^W^*/*dy^W^*muscle. Importantly, AAV9-Spc/CBh-DL-injected healthy control mice showed only minimal transcriptional changes (Figure S6B). We next compared the transcriptome of untreated *dy^W^*/*dy^W^* mice with untreated wild-type mice using a threshold for differentially expressed genes of log2±0.5 and an adjusted p ≤ 0.05. Out of the 14,905 detected transcripts, 14% were significantly downregulated and 18% were significantly upregulated (Figure 6A). Among the most significantly upregulated genes were *Sln* and *Cilp*, encoding sarcolipin and cartilage intermediate layer protein, respectively – both known markers of dystrophic pathology.^39,40^ Consistent with the strong ameliorative effect of the two linker proteins on the pathology in *dy^W^*/*dy^W^*mice, expression of the dysregulated genes was changed towards wild-type in *dy^W^*/*dy^W^* AAV9-Spc/CBh-DL mice (Figure 6B), with their transcriptome being closer to wild-type controls than untreated *dy^W^*/*dy^W^*mice (Figure S6A). This “normalization” of the gene expression profiles was clearly confirmed by the very strong correlation (R^2^ > 0.6) between genes changed in *dy^W^*/*dy^W^* mice (log2 (*dy^W^*/*dy^W^*/wild-type) and genes changed in *dy^W^*/*dy^W^* mice after treatment (log2 (*dy^W^*/*dy^W^* AAV9-Spc/CBh-DL/ *dy^W^*/*dy^W^*) (Figure S6C).

**Figure 6.**
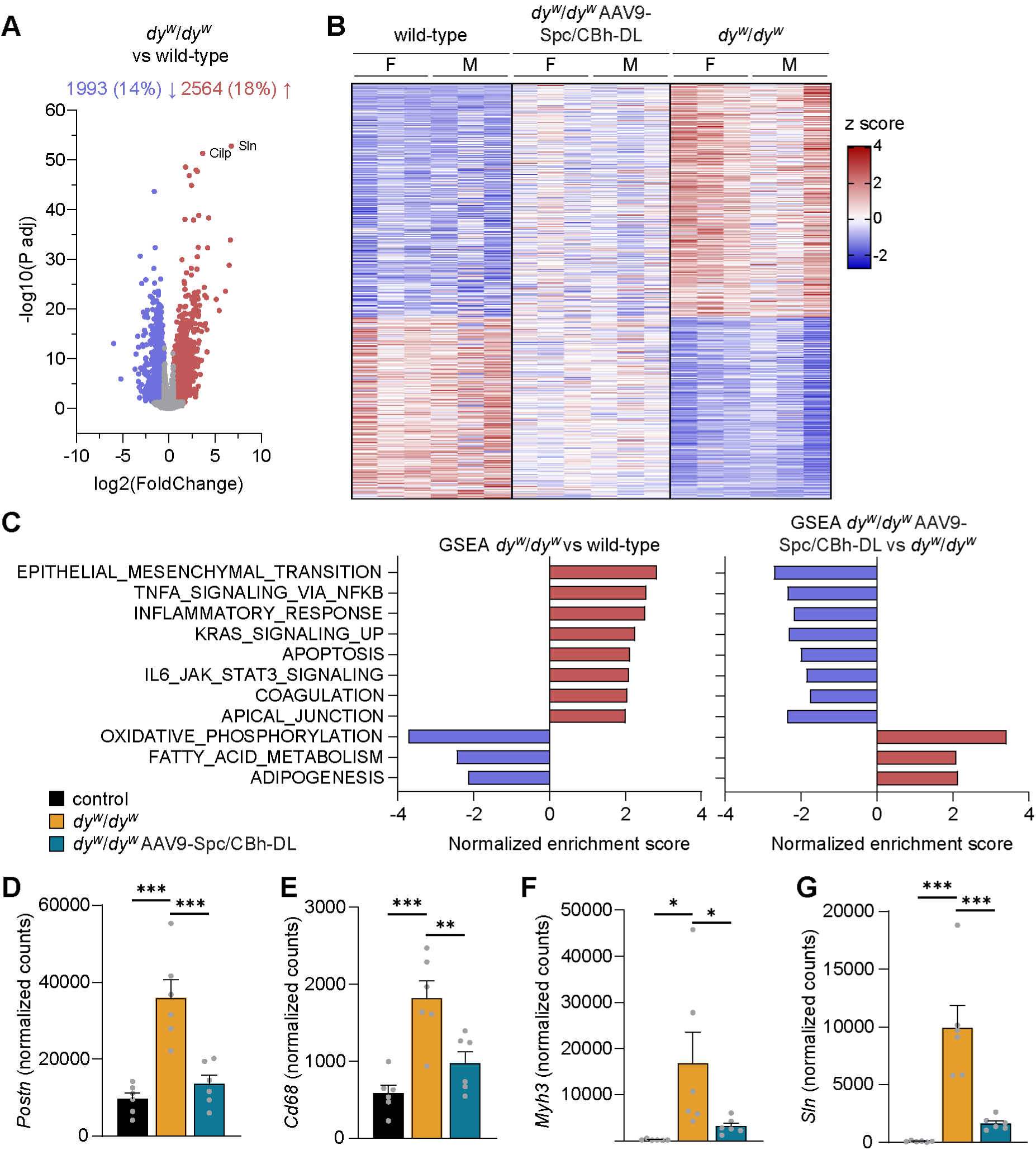
Normalization of dysregulated transcriptional hallmarks by ubiquitous expression of αLNNdΔG2 in combination with muscle-specific expression of mag. (**A-G**) RNAseq analysis of TA muscle from 8-week-old wild-type, *dy^W^*/*dy^W^* and *dy^W^*/*dy^W^* mice injected intravenously at P1 with AAV9-Spc/CBh-DL (1×10¹⁴ vg/kg per construct). (**A**) Differentially expressed transcripts in *dy^W^*/*dy^W^* versus wild-type mice presented as a Volcano plot. Differentially expressed genes (DEGs) are shown in red (upregulated) or blue (downregulated), based on a threshold of log2 fold change > ±0.5 and adjusted p < 0.05. Numbers on top indicate number and percentage of upregulated (red) or downregulated (blue) transcripts. (**B**) Heatmap of z-scores for the 4,557 DEGs in *dy^W^*/*dy^W^* versus wild-type mice, and corresponding *z*-scores in *dy^W^*/*dy^W^* AAV9-Spc/CBh-DL mice. (**C**) (Left) Gene set enrichment analysis (GSEA) of Hallmark Gene Set (MSigDB) comparing *dy^W^*/*dy^W^* and wild-type muscle. Gene sets with a normalized enrichment score (NES) > ±2 are shown. (Right) NES values for the same gene sets in *dy^W^*/*dy^W^* AAV9-Spc/CBh-DL versus untreated *dy^W^*/*dy^W^* mice. (**D-G**) Normalized transcript counts for selected genes: (**D**) *Postn* (marker of fibrosis) (**E**) *Cd68* (marker for inflammation) (**F**) *Myh3* (marker for fiber degeneration/regeneration) (**G**) *Sln* (associated with calcium homeostasis). Data are presented as mean ± SEM. \**P* < 0.05, \*\**P* < 0.01, \*\*\**P* < 0.001 by one-way ANOVA with Dunnett’s post hoc test, comparing each group to untreated *dy^W^*/*dy^W^*mice. *n* = 6 mice per group.

Gene set enrichment analysis ^41^ (MSigDB^35^) identified hallmark pathways changed in *dy^W^*/*dy^W^* muscle, including increases in transcripts involved in fibrosis (epithelial-mesenchymal transition, TNF-α signaling), inflammation and apoptosis, alongside downregulated transcripts associated with oxidative phosphorylation, fatty acid metabolism, and adipogenesis. Notably, all hallmark pathways responded positively to treatment (Figure 6B). Fibrosis and inflammation markers (*Postn*, *Cd68*), previously shown to be highly upregulated in laminin-α2 deficiency,^42–44^ were significantly reduced post-treatment (Figure 6D-E). *Myh3*, an indicator of active muscle regeneration, was also significantly downregulated, suggesting reduced degeneration/regeneration cycles despite persistent central nucleation (Figure 6F). *Sln*, the most strongly dysregulated gene in *dy^W^*/*dy^W^* mice, also responded robustly to treatment (Figure 6G). All-in-all, these results are further support for the effectiveness of the linker proteins to normalize the disease signature towards wild-type mice.

### Functional measures reach near-normal values in *dy^W^*/*dy^W^* mice treated early with AAV9- or AAV8-delivering muscle-expressed mag and ubiquitously-expressed αLNNdΔG2

During our studies, we observed that some *dy^W^*/*dy^W^*mice treated at P1 with AAV9-Spc/CBh-DL were nearly indistinguishable from their healthy littermates at 8 weeks of age (Video S5, Figure 7A). To evaluate this observation quantitatively, we subjected the mice to a range of functional assessments. As observed with muscle-specific expression of the linker proteins (Figure 2D), body weight in treated *dy^W^*/*dy^W^* mice remained comparable to that of untreated *dy^W^*/*dy^W^* mice until approximately 3 weeks of age. However, beyond this time point, AAV9-Spc/CBh-DL-injected *dy^W^*/*dy^W^* mice began to gain weight at a similar rate as their control littermates (Figure 7B). At 8 weeks of age, gait speed, a readout of peripheral nerve function, was restored to near-normal levels in treated *dy^W^*/*dy^W^*mice (Figure 7C). Furthermore, when normalized to body mass, muscle mass in AAV9-Spc/CBh-DL-treated *dy^W^*/*dy^W^* mice were very similar to those of healthy controls (Figure S7A).

**Figure 7.**
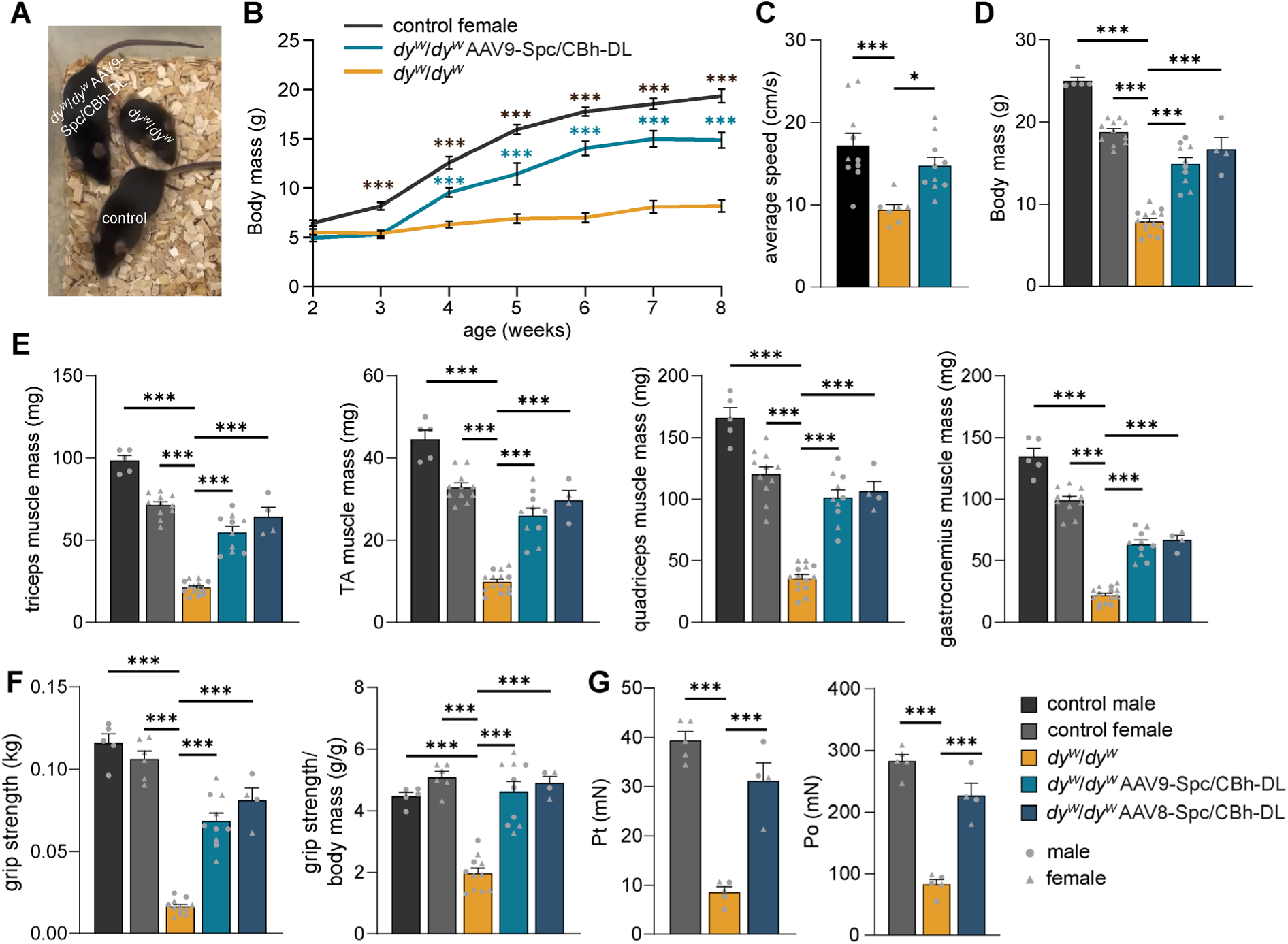
Functional improvement of AAV9-or AAV8-mediated dual expression of mag in muscle and αLNNdΔG2 in muscle and nerve in *dy^W^*/*dy^W^* mice. (**A**) Picture of representative 8-week-old healthy control, untreated *dy^W^*/*dy^W^,* and *dy^W^*/*dy^W^* mice injected at P1 with AAV9-Spc/CBh-DL (1×10¹⁴ vg/kg per construct). (**B**) Body mass curves from 2 to 8 weeks of age. (**C**) Quantitative analysis of locomotor speed at 8 weeks of age using a gait analysis system. (**D-G**) Analysis in 8-week-old healthy control, untreated *dy^W^*/*dy^W^,* and *dy^W^*/*dy^W^* mice injected at P1 with AAV9-Spc/CBh-DL or AAV8-Spc/CBh-DL (1×10¹⁴ vg/kg per construct). (**D**) Comparison of body mass at 8 weeks of age. (**E**) Comparison of the mass of fore- and hindlimb muscles from 8-week-old mice. (**F**) Absolute and body mass-normalized forelimb grip strength of 8-week-old mice. (**G**) Absolute twitch (Pt) and tetanic force (Po) measurements of isolated extensor digitorum longus (EDL) muscles. Data are presented as mean ± SEM. \**P* < 0.05, \*\**P* < 0.01, \*\*\**P* < 0.001 by one-way ANOVA with Dunnett’s post hoc test, comparing each group to untreated *dy^W^*/*dy^W^* mice. *n* = 5-14 mice per group.

Given the favorable safety and efficacy profile of AAV8 in recent clinical trials,^45,46^ we also tested AAV8 as a delivery vehicle for the linker proteins. To compare targeting efficiency between AAV8 and AAV9, we used a CMV-GFP construct packaged into either serotype.

Consistent with previous reports,^47^ muscle transduction was comparable between AAV8 and AAV9 (Figure S7B). Importantly, AAV8 also efficiently transduced peripheral nerves (Figure S7C). Indeed, the therapeutic efficacy of AAV8-Spc/CBh-DL in *dy^W^*/*dy^W^*mice mirrored that observed with AAV9. Body mass (Figure 7D) and muscle mass (Figure 7E) at 8 weeks of age were indistinguishable from those achieved with AAV9-mediated delivery. Functional assessments revealed improved grip strength (Figure 7F) and *ex vivo* EDL force generation (Figure 7G), both of which are severely impaired in untreated *dy^W^*/*dy^W^* mice. In line with these functional and weight improvements, muscle histology was similarly improved with AAV8 (Figure S8A), as with AAV9 treatment (Figure 5), reflected by reduced fibrosis (Figure S8B), increased muscle fiber number (Figure S8C), and normalization of fiber size distribution (Figure S8D-E). Taken together, these results demonstrate that neonatal administration of Spc/CBh-DL via either AAV8 or AAV9 restores near-normal function and structure in the muscle and the peripheral nerve of *dy^W^*/*dy^W^* mice by 8 weeks of age.

### Late treatment still improves defects in *dy^W^*/*dy^W^* mice

In patients with severe LAMA2 MD, symptoms are present at birth or become evident by six months of age.^6^ Similarly, *dy^W^*/*dy^W^*mice exhibit an early-onset, severe phenotype that progressively worsens over time. To determine if the linker treatment can improve the disease at more advanced disease stages, we assessed the treatment effect of the linkers when administered at three weeks of age. At this stage, *dy^W^*/*dy^W^* mice show significant muscle impairment, pronounced fibrosis, and clear signs of peripheral neuropathy. This time point also coincides with the onset of increased mortality in untreated *dy^W^*/*dy^W^* mice. To accommodate the higher body weight and slower growth rate of older mice - and to remain within the upper bounds of vector capacity while achieving therapeutic saturation - we intravenously injected 5×10¹³ vg/kg of each vector. This dose aligns with those safely administered in clinical trials for neuromuscular diseases.^45^ To quantitatively assess therapeutic efficacy, grip strength was measured at 8 weeks of age, followed by rotarod performance and TA muscle histology at 12 weeks (Figure 8A).

**Figure 8.**
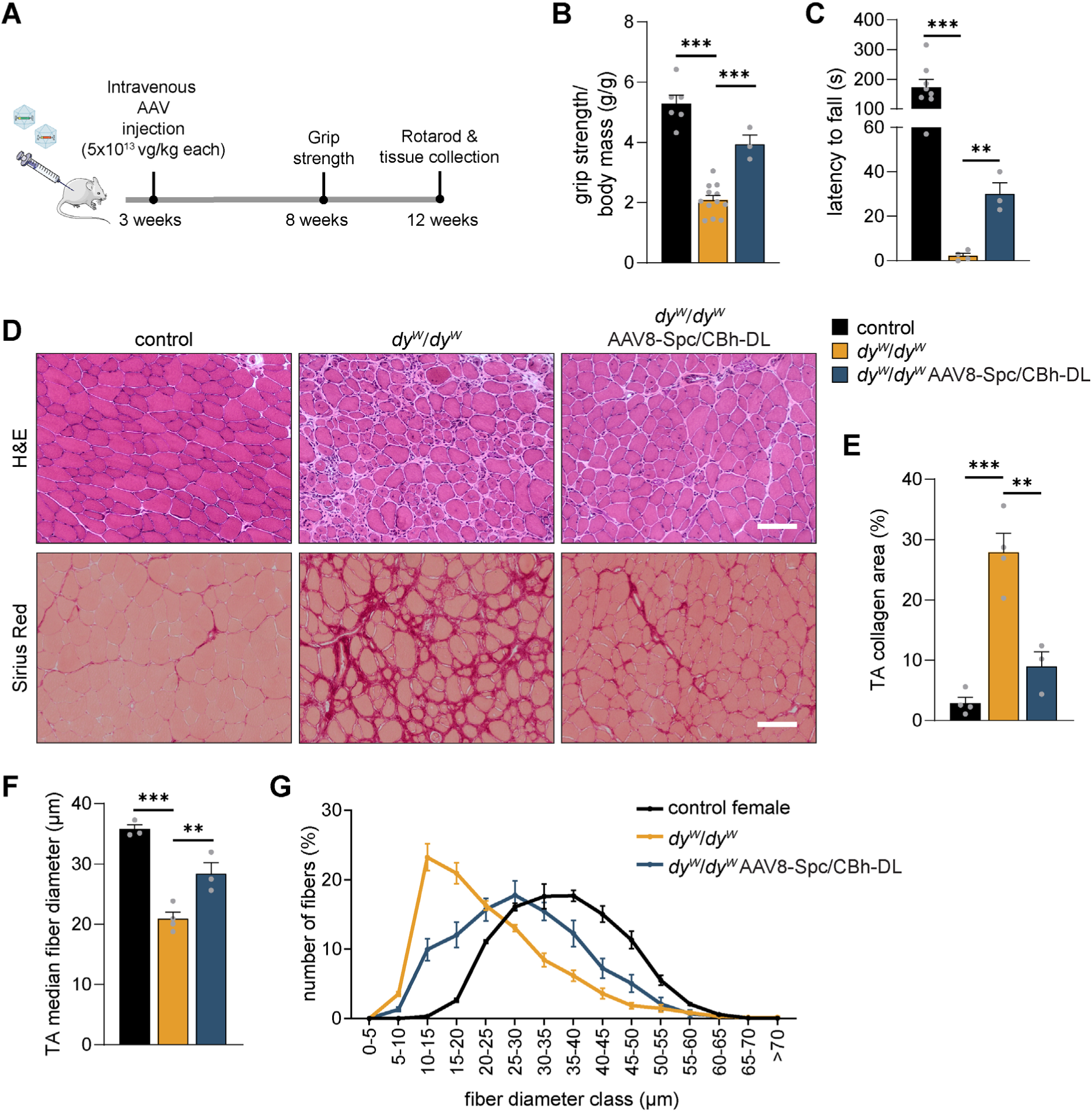
Therapeutic effects of late AAV8-mediated expression of mag in muscle and αLNNdΔG2 in muscle and nerve on muscle histology and neuromuscular function in *dy^W^*/*dy^W^* mice. (**A**) Experimental design for panels B-G: *dy^W^*/*dy^W^* mice were injected intravenously at 3 weeks of age with AAV8-Spc/CBh-DL (5×10^13^ vg/kg per construct). Forelimb grip strength was analyzed at 8 weeks, rotarod performance at 12 weeks, and tissues were collected at 12 weeks. (**B**) Forelimb grip strength normalized to body mass. (**C**) Rotarod-based assessment of motor function and coordination. (**D**) Representative images of H&E- and Sirius Red-stained TA muscle cross-sections. (**E**) Quantification of fibrosis based on Sirius Red-positive area in TA muscle cross-sections. (**F**) Quantification of median fiber diameters in TA cross-sections. (**G**) Distribution of fiber diameters in TA cross-sections. Data are presented as mean ± SEM. \**P* < 0.05, \*\**P* < 0.01, \*\*\**P* < 0.001 by one-way ANOVA with Dunnett’s post hoc test, comparing each group to untreated *dy^W^*/*dy^W^* mice. Scale bar: 100 μm. *n* = 3-12 mice per group.

AAV8-Spc/CBh-DL-treated *dy^W^*/*dy^W^* mice exhibited a markedly improved phenotype, with particularly notable effects on the progression of peripheral neuropathy. Hindlimb dragging, a common feature in untreated 12-week-old *dy^W^*/*dy^W^* mice, was noticeably reduced in treated animals (Movie S6). In grip strength analysis, untreated *dy^W^*/*dy^W^* mice that reached 12 weeks of age showed a further decline in muscle function compared to those at 8 weeks; in contrast, treated mice were significantly improved (Figure 8B). Additionally, the severe peripheral neuropathy in untreated mice limited their ability to stay on a rotating rod, with latency to fall lasting only a few seconds. Treated mice, however, exhibited a significantly increased latency to fall, reflecting enhanced motor coordination and strength (Figure 8C).

Histological analysis of the TA muscle of 12-week-old mice revealed clear histological improvement in treated *dy^W^*/*dy^W^* mice (Figure 8D). Notably, fibrosis was strikingly reduced, as evidenced by the significant decrease in collagen-positive area in Sirius Red-stained sections (Figure 8E). Furthermore, the median muscle fiber diameter was increased (Figure 8F), and the fiber size distribution shifted toward larger fibers (Figure 8G). In summary, these results demonstrate that AAV8-Spc/CBh-DL treatment not only ameliorates the disease phenotype when administered neonatally but also provides substantial therapeutic benefit when delivered at a later stage, when both muscular dystrophy and peripheral neuropathy are advanced.

## DISCUSSION

Gene replacement therapy using AAV vectors is particularly challenging for LAMA2 MD, primarily due to the large size of the *LAMA2* coding sequence, by far exceeding the packaging capacity of AAVs. Additionally, immune responses against transgene products pose a significant concern, not only threatening long-term expression but also raising serious safety issues, as recently demonstrated in gene therapy trials for Duchenne muscular dystrophy (DMD). ^14^ Beyond the anti-transgene immune response, anti-capsid immunity represents a second major safety challenge in human applications. ^48^ Reports from clinical studies have described cases of thrombotic microangiopathy resulting from complement activation, particularly following administration of AAV9 or novel capsids.^49,50^. To address this concern, we evaluated AAV8, for which no such adverse events have been reported to date. Our goal was to evaluate an AAV capsid with the most favorable safety profile for future human use.

In this study, we demonstrate therapeutic efficacy on both muscle and nerve pathology in the severe LAMA2 MD mouse model by using two engineered linker proteins derived from ECM components naturally found in patients. This strategy leverages the presence of laminin-4X1—an alternative laminin isoform that is consistently upregulated in LAMA2 MD but lacks sufficient receptor binding and polymerization capacity to fully compensate for the loss of laminin-2X1.^21^. We have previously shown that disease progression can be mitigated by transgenic expression of mini-agrin (mag)^51–53^ and further improved through the addition of a second linker protein that restores polymerization.^21^ Here, we optimized these linker proteins for effective AAV-mediated delivery, with a view toward clinical translation.

Importantly, we show that muscle-restricted expression of the linker proteins improves muscular dystrophy with long-term benefits. By employing myotropic AAVs, we achieved therapeutic efficacy at lower vector doses. However, this muscle-targeted strategy was insufficient to prevent progressive neuropathy, a hallmark of LAMA2 MD. Although peripheral neuropathy is well-documented in LAMA2 MD mouse models, its clinical relevance in human patients remains less clearly defined.^12^ Nerve conduction abnormalities have been reported in several patients, ^54^ and severe peripheral neuropathy has been described particularly in those with late-onset LAMA2 MD.^55–57^ Our findings in mice highlight that while neuropathy appears mild at early stages, it progresses significantly over time—particularly when survival is extended by treating muscle pathology. This suggests that neuropathy may similarly progress in human patients unless therapeutic approaches address both muscle and nerve deficiencies.

We previously demonstrated that combined transgenic expression of both linker proteins improves nerve pathology in *Lama2*-null mice.^31^ To reduce reliance on ubiquitous promoters, we now limit their use to settings where direct nerve targeting is essential. Because the mechanisms underlying nerve pathology are less well understood^12^ than those affecting muscle, we generated mouse models with ubiquitous expression of each linker protein individually to dissect their respective contributions. Our data show that αLNNd alone is sufficient to ameliorate nerve pathology. In contrast, mice expressing mag in peripheral nerves still developed hindlimb paralysis, consistent with earlier findings,^58^ indicating that laminin polymerization plays a particularly important role in preventing nerve pathology.

Supporting this, previous studies have shown that peripheral neuropathy in laminin-α2-null mice can be rescued by transgenic expression of polymerizing laminin-α1.^59^. Additionally, in models expressing truncated laminin-α2, restoration of laminin polymerization - either through a linker protein^36^ or via dCas9-mediated upregulation of *Lama1*^60^ - has also been shown to improve nerve pathology.

Based on these insights, we proceeded with a combination therapy consisting of muscle-specific mag and ubiquitously expressed αLNNd. To enable efficient expression, αLNNd was shortened and placed under the control of the CBh promoter. When this dual therapy was delivered neonatally via AAV8 or AAV9 at clinically relevant doses, treated *dy^W^*/*dy^W^* mice achieved near-normal body weight, histology, and functional outcomes.

To evaluate the global effect of treatment, we performed unbiased transcriptomic analysis, revealing normalization of gene expression across major disease-related pathways. Key dysregulated processes in LAMA2 MD muscle - including fibrosis,^61^, inflammation,^42^ apoptosis,^62^ and reduced oxidative phosphorylation^63^ - were all largely corrected by the linker protein treatment. Many of these pathways are commonly implicated in other forms of muscular dystrophy, and their targeting may yield functional benefits. However, unlike interventions targeting these secondary effects, the linker protein-based therapy addresses the root cause of the disease, leading to broader correction. Moreover, the transcriptomic data may serve as a valuable resource for identifying biomarkers to monitor therapeutic efficacy in future clinical trials.

The post-mitotic nature of muscle fibers is favorable for the long-term maintenance of AAV-delivered episomes. However, in the context of muscular dystrophy, increased muscle turnover and regeneration may dilute expression over time. In this regard, the secreted nature of the linker proteins offers a key advantage in that newly formed fibers may still benefit from linkers secreted by neighboring transduced cells. While dual-AAV approaches present challenges for cost-effective manufacturing, each individual linker confers therapeutic benefit. Due to their extracellular mode of action, both linkers need not be co-expressed in the same cell- unlike recombination- or intein-based strategies that require co- localization within single fibers.

Although laminin-2X1 surrounds muscle fibers, recent single-cell RNA-sequencing studies suggest that muscle fibers may not be the primary source of laminin-2X1.^64–66^ This complicates gene editing strategies, which rely on targeting the correct cell type. In contrast, our linker proteins are captured by laminin-4X1 at the muscle fiber basement membrane and do not need to be expressed by the laminin-producing cells themselves.

While neonatal treatment restored near-normal muscle and nerve function, treatment at later stages was less effective - though still conferred substantial therapeutic benefit. This reduced efficacy may be attributed to extensive fibrotic tissue accumulation, impaired muscle regeneration in advanced stages of LAMA2 MD^67^ or disruptions in integrin signaling^68^ aspects not yet fully explored in the context of linker protein therapy.

Consistent with previous reports, we observed no reduction in the proportion of muscle fibers with centralized nuclei in treated animals.^21,31^ Interestingly, our RNA-seq data showed a marked downregulation of gene expression markers associated with active regeneration. This discrepancy suggests that centralized myonuclei in treated mice may reflect earlier degeneration-regeneration events, rather than an ongoing pathology - similar to observations following acute muscle injury.^69,70^ Recent findings in other dystrophies and myopathies also reported the persistence of centralized nuclei despite clear functional improvements after treatment.^71,72^ This may be explained by the initiation of therapy at a stage when substantial pathology is already present.

In summary, this study addresses key limitations of AAV-based gene therapy for LAMA2 MD by engineering linker proteins derived from domains of other ECM components to replace the function of the missing laminin-α2 chain. These proteins can be efficiently delivered via AAV and act extracellularly, making them compatible with the constraints of AAV vector size and distribution. This strategy not only provides a practical alternative to full gene replacement but also offers an example for treating other diseases caused by mutations in large genes that exceed AAV packaging limits.

## MATERIALS AND METHODS

### Mice

As a mouse models for LAMA2 MD, we used *dy^W^*/*dy^W^* mice [B6.129S1(Cg)- Lama2tm1Eeng/J; available from the Jackson Laboratory stock #013786].^32,73^ Genotyping was performed as previously described.^32^

Transgenic knock-in mag mice [C57BL/6J-Gt(ROSA)26Sortm1(CAG-mag)Rueg] and αLNNd mice [C57BL/6J-Gt(ROSA)26Sortm2(CAG-aLNNd)Rueg] were described previously.^31^ To achieve ubiquitous expression of the transgenes, these lines were crossed with CAG-Cre-ER^TM^ mice [B6.Cg-Tg(CAG-cre/Esr1*)5Amc/J ^74^ available from the Jackson Laboratory stock 004682]. As reported previously,^31^ recombination and transgene expression occurred in muscle and peripheral nerve tissue even in the absence of tamoxifen, and thus no tamoxifen was administered in this study.

All experimental animals were obtained from breedings of mice heterozygous for the *Lama2* knockout allele, enabling the generation of all genotypes within a single litter and the use of littermate controls. Control animals were either wild-type or heterozygous for the *Lama2* locus. For the transgenic linker protein-expressing lines, controls were always Cre-negative and either heterozygous or homozygous for the LSL-αLNNd and LSL-mag alleles. Unless otherwise indicated, both male and female mice were used. To support the dystrophic mice, all cages were equipped with long-necked water bottles and provided with wet food from weaning onward.

All animal experiments were conducted in accordance with federal guidelines and approved by the Veterinary Authority of the Canton of Basel-Stadt.

### AAV vector cloning, production, purification and titration

Mag and αLNNd constructs were subcloned into AAV plasmids containing either the Spc5.12 (Spc)^27,28^ or the CBh promoter^37^. Expression cassettes were flanked by AAV2 inverted terminal repeats (ITRs) and included either a bovine growth hormone (BGH) polyA for mag and αLNNdΔG2, or a synthetic small polyA ^75^ for full-length αLNNd.

AAV vectors were produced by triple transfection of adherent HEK293 cells and purified by iodixanol gradient ultracentrifugation. AAVs were obtained either from commercial providers (Vigene Biosciences or Charles River Laboratories) or produced in-house as previously described.^76^ The following capsid and helper plasmids were used: AAV8 and AAV9 (gifts from J.M. Wilson; Addgene plasmids #112864 and #112867), AAVMYO^30^ (gift from D. Grimm), AAVMYO ^38^ (gift from D. Grimm), and pAdDeltaF6 (gift from J.M. Wilson; Addgene plasmid #112867). Purity and particle concentration of AAV preparations were assessed by SDS-PAGE followed by Coomassie Blue staining.

For AAV titration, unencapsidated DNA was digested with DNase I (New England Biolabs [NEB], M0303). Next, samples were serially diluted in dilution buffer (10 mM Tris-HCl (pH 8.0), 0.1 mM EDTA, 100 μg/ml poly(A), and 0.01% Pluronic F-68. Capsids were lysed by heating to 95 °C for 10 minutes, and viral DNA was linearized using MspI (NEB, R0106).

TaqMan®-based digital PCR was performed on QIAcuity Nanoplates (26k 24-well, Qiagen) using the QIAcuity One 5plex system (Qiagen). Primer/probe sets targeted the ITR (dPCR CGT Assay ITR2/5, FAM; Cat. No./ID: 250230), BGH polyA (HEX; Cat. No./ID: 250234), and either mag (forward: 5′-GGC AGA ACC TTC GTG GAG TA; reverse: 5′-TCT CAG GCT CAG CTC GAA GT; probe: 5′-ACG CCG TGA CCG AGA GCG AG, Cy5-labeled) or αLNNd (forward: 5′-CAC CAC CAT CAT CAG ACA GG; reverse: 5′-GCT GTC GGT CCA GAA GAT GT; probe: 5′-CGT GGA CCA CCT GGG CAG AA, Cy5-labeled).

### AAV administration

For neonatal injections, a single intravenous injection was performed on postnatal day 1 via the temporal facial vein using a 30G insulin syringe. AAV vectors were diluted in phosphate-buffered saline (PBS) and administered at a volume of 50 μl per pup. After injection, pups were placed on a heating pad and monitored until recovered before being returned to the dam.

For injections at 3 weeks of age, mice received an intravenous injection into the lateral tail vein. A total dose of 1×10¹⁴ vg/kg was administered at a volume of 20 μl per gram of body weight.

### AAV genome copy number quantification

Tissues were pulverized in liquid nitrogen and genomic DNA was extracted using the DNeasy® Blood & Tissue Kit (Qiagen) according to the manufacturer’s instructions. Viral genome copy numbers per diploid mouse genome were determined by quantitative PCR (qPCR) using a QuantStudio 5 Real-Time PCR System (Applied Biosystems) and PowerUp SYBR Green Master Mix (Applied Biosystems). Specific primers were used to amplify mag (mag_f: 5′-TGT ACA ACG GCC AGA AGA CC; mag_r: 5′-TCT CTG CTT CTG ATC ACG GC), αLNNd (αLNNd_f: 5′-ACA GAA TCG AGG TGG CCA AG; αLNNd_r: 5′-GGT CCA GTA CAG GTT GCC TC), and the mouse *Rosa26* locus (Rosa26_f: 5′-GTG GAG CCG TTC TGT GAG AC; Rosa26_r: 5′-CTT TTC CGC TCC CTT CTC CC). Copy numbers were calculated using standard curves generated from serial dilutions of the corresponding transfer plasmids and the Ai9 plasmid (a gift from H. Zeng Addgene plasmid # 22799) as a reference for the *Rosa26* locus.

### Antibodies

The following antibodies were used for immunostaining and Western blot analyses: agrin (for detection of m.mag; produced in-house ^77^; αLNNd (for detection of αLNNd by immunostainings, previously described ^78^; 1:100); F4/80 (Bio-Rad, MCA497G, clone CI:A3-1; 1:50); GFP (Molecular Probes A10262; 1:400); laminin-α1 (for detection of αLNNd; R&D Systems, AF4187; 1:2,000 for Western blot, 1:200 for immunostaining); laminin-α2 (Sigma-Aldrich, L0663, clone 4H8-2; 1:500); laminin-α4 (as previously described^78^; 1:1,000 for Western blot, 1:200 for immunostaining); laminin-β1γ1 (Sigma-Aldrich, L9393; 1:100); and GAPDH (Cell Signaling Technology, 2118; 1:1,000).

Antibodies against the human version of mag were generated in-house. The mag sequence was cloned into the pCEP expression vector containing a puromycin resistance cassette and a 6×His tag. HEK293 cells were transfected using Lipofectamine 2000 (Invitrogen) according to the manufacturer’s instructions and cultured in DMEM/F12 medium (Invitrogen) supplemented with 10% fetal calf serum. Two days post-transfection, medium was replaced with selection medium containing 2 μg/ml puromycin (Sigma-Aldrich), and stable clones were expanded. Upon reaching 80–90% confluence, cells were switched to serum-free DMEM/F12. Conditioned medium was collected after 2-3 days and protein production was validated by SDS-PAGE followed by Coomassie Blue staining.

For purification, 10× supernatant buffer (500 mM NaH₂PO₄, 1.5 M NaCl, 100 mM imidazole, pH 8.0) was added to the conditioned medium, which was then incubated overnight at 4 °C with rotation using pre-washed Ni-NTA agarose (Qiagen). The resin was washed four times with wash buffer (50 mM NaH₂PO₄, 300 mM NaCl, 20 mM imidazole, pH 8.0), and protein was eluted with elution buffer (50 mM NaH₂PO₄, 300 mM NaCl, 250 mM imidazole, pH 8.0). The eluate was dialyzed against PBS and purified mag-6×His was validated by SDS-PAGE and Coomassie Blue staining. Protein concentration was determined using a BCA assay.

Rabbit polyclonal antibodies were generated by immunizing two rabbits with purified mag-6×His (Davids Biotechnologie GmbH). Pre-immune serum from five rabbits was screened by Western blot analysis and immunostaining to identify cross-reactivity. Two rabbits were selected and received five immunizations (days 1, 14, 28, 42, and 56). Test serum was collected on day 35 and final serum on day 63. Non-purified antiserum was used for Western blot analysis and immunostaining at a dilution of 1:20,000.

### Immunostainings

Muscle and sciatic nerve tissues were embedded in Tissue-Tek® O.C.T. compound (Sakura) and rapidly frozen in 2-methylbutane pre-cooled in liquid nitrogen (-150°C). For GFP analysis, tissues were fixed immediately after dissection in ice-cold 4% paraformaldehyde (PFA) for 2 h at 4°C, followed by dehydration in 30% sucrose overnight at 4°C, then embedded in O.C.T. and frozen. Frozen tissues were cryosectioned at a thickness of 10 μm and either fixed in 4% PFA in PBS or processed unfixed.

Sections were blocked for 1 h at room temperature in PBS containing 5% donkey serum and 0.3% Triton X-100. Primary antibodies were diluted in 5% donkey serum in PBS and incubated overnight at 4°C. For stainings involving mouse or rat primary antibodies, endogenous mouse IgG was blocked using M.O.M. Blocking Reagent (Vector Laboratories, MKB-2213; 1:40) added to the blocking buffer. Following three PBS washes, sections were incubated with appropriate secondary antibodies for 1 h at room temperature, washed again, and mounted using ProLong™ Gold Antifade Mountant (Invitrogen). Imaging was performed using an Olympus iX81 microscope, a Zeiss Axio Scan.Z1 slide scanner, or a Zeiss LSM700 confocal laser scanning microscope.

### Histology and histological quantifications

Muscles were embedded in Tissue-Tek® O.C.T. compound (Sakura) and rapidly frozen in 2- methylbutane cooled in liquid nitrogen (-150°C). Transverse sections (10 μm thickness) were cut on a cryostat. For general histological analysis, sections were fixed in 4% paraformaldehyde and stained with hematoxylin and eosin (Merck) or Picro Sirius Red (Direct Red 80; Sigma) in picric acid solution. Images were acquired using an Olympus iX81 microscope equipped with cellSens software (Olympus).

Muscle fiber diameter was quantified using the minimal Feret’s diameter (the shortest distance between parallel tangents at opposing borders of the fiber), as previously described^79^. For analysis of fiber number, fiber size, and centrally nucleated fibers (CNFs), complete mid-belly cross-sections were evaluated using an in-house customized, Fiji-based version of Myosoft.^31,80^ Muscle fibrosis was quantified by measuring hydroxyproline content via mass spectrometry-based amino acid analysis. Frozen muscles were pulverized in liquid nitrogen and dried using a SpeedVac. Quantification was performed as described previously. ^31^ For histological analysis of sciatic nerves, nerves were dissected and fixed in 2.5% glutaraldehyde in 0.12 M phosphate buffer (pH 7.4), post-fixed with 1% osmium tetroxide, embedded in Epon, and processed as previously described.^31^

### Protein extraction and Western blot analysis

For total protein extracts, frozen muscle or sciatic nerve tissue was pulverized in liquid nitrogen and lysed in modified RIPA buffer (50 mM Tris-HCl pH 8.0, 150 mM NaCl, 1% NP- 40, 0.5% sodium deoxycholate, 0.1% SDS, 20 mM EDTA, supplemented with protease inhibitors). Lysates were sonicated and incubated for 2 hours at 4 °C with end-over-end rotation. Insoluble material was removed by centrifugation at 16,000 × g for 30 minutes at 4 °C. Protein concentrations were determined using the BCA assay, and samples were adjusted accordingly. Laemmli buffer was added, and samples were denatured by heating for 5 minutes at 95 °C before subjecting them to standard SDS-PAGE and Western blot analysis.

### LC-MS Analysis

Frozen muscle tissues were pulverized in liquid nitrogen and resuspended in lysis buffer (5% SDS, 10mM TCEP, 0.1 M TEAB), followed by sonication using a PIXUL Multi-Sample Sonicator (Active Motif). Lysates were incubated at 95°C for 10 min and subsequently alkylated by 20 mM iodoacetamide at 25°C for 30 min. For each sample, 50 µg protein was digested using S-Trap™ micro spin columns (Protifi) according to the manufacturer’s instructions, employing sequencing-grade modified trypsin (1/25, w/w; Promega). After elution, peptides were dried under vacuum. Dried peptides were resuspended in 0.1% aqueous formic acid and subjected to LC-MS/MS analysis using an Orbitrap Eclipse Tribrid Mass Spectrometer fitted with Ultimate 3000 nano system (Thermo Fisher Scientific) and a custom-made column heater set to 60°C. Peptides were resolved using a RP-HPLC column (75 μm × 30 cm) packed in-house with C18 resin (ReproSil-Pur C18–AQ, 1.9 μm resin; Dr. Maisch GmbH) at a flow rate of 0.3 μL/min. The following gradient was used for peptide separation: from 2% B to 12% B over 5 min, to 35% B over 45 min, to 50% B over 10 min, to 95% B over 2 min, followed by 18 min at 95% B, then back to 2% B over 2 min, followed by 18 min at 2% B. Buffer A was 0.1% formic acid in water and buffer B was 80% acetonitrile, 0.1% formic acid in water.

The mass spectrometer was operated in DIA mode with a cycle time of 3 seconds between master scans. MS1 spectra were acquired in the Orbitrap at a resolution of 120,000 and a scan range of 390 to 920 m/z, AGC target set to 1E6 and maximum injection time set to 45 ms. A total of 46 DIA windows with an isolation window size of 11 m/z and a window overlap of 1 m/z were recorded. Precursor mass range was set to 400-900 m/z, AGC target to 5E5 and a maximum injection time to 22 ms. The acquired files were searched using the Spectronaut (Biognosys version 17.4.230317.55965) directDIA workflow against the Mus musculus protein database (UniProt, download Feb 22nd, 2022) with the addition of human proteins agrin, nidogen-1 and laminin-α1. Quantitative fragment ion data (F.Area) was exported from Spectronaut and analyzed using the MSstats R package v.4.8.0. (https://doi.org/10.1093/bioinformatics/btu305). Data was normalized using the default normalisation option “equalizedMedians”, imputed using the “AFT model-based imputation”.

### RNA-seq analysis

Tibialis anterior (TA) muscles were pulverized on a metal block cooled with liquid nitrogen. Total RNA was extracted using the RNeasy Fibrous Tissue Mini Kit (Qiagen) following the manufacturer’s instructions. RNA concentration was quantified using the Qubit RNA HS assay (Thermo Fisher Scientific), and RNA integrity was assessed on an Agilent TapeStation. Libraries were prepared using the TruSeq RNA Library Prep Kit v2 (Illumina) starting from 200 ng of RNA. RNA sequencing was performed using the Illumina NovaSeq 6000 system (PE 2 X 51), producing 150-bp paired-end reads. Sequencing resulted in 50-55 million reads per sample.

Raw transcript reads were aligned to the *Mus musculus* reference genome (GRCm38/mm10) using Salmon (v.1.1.0) with the flags validateMappings, seqBias, and gcBias. Normalization and differential expression analysis were completed using the R package DESeq2 (Bioconductor). Transcript level information was summarized to gene level. Initial filtering was applied by retaining only genes with a raw count >10 in at least three samples per group. Genes were kept only if this criterion was met in all groups.

Unsupervised sample clustering was performed using correlation distance as the distance metric and complete linkage for hierarchical clustering. Pairwise sample-to-sample correlations were visualized as a heatmap using pheatmap R package with annotated sample conditions.

### Grip strength measurement

Forelimb grip strength was assessed in accordance to SOP MDC1A_M.2.2.001 (TREAT-NMD) using a grip strength meter (Columbus Instruments) equipped with a trapeze bar. Mice were lifted by the tail and guided toward the bar, allowing them to grasp it with both forepaws. Once a secure grip was established, the mouse was gently pulled away at a constant speed until it released the bar. The peak force was recorded, and the final force was calculated as the mean of the three highest values from six consecutive trials.

### Gait analysis

Gait analysis was performed using the CatWalk XT system (Noldus) in accordance with the manufacturer’s instructions. Mice were placed in an enclosed, illuminated glass walkway and allowed to move freely in both directions. Paw placement was recorded by a high-speed video camera positioned beneath the walkway. Runs were considered compliant if the mouse traversed a defined 40 cm segment within 10 seconds and maintained a maximum speed variation of ≤60%. For each mouse, three compliant runs were recorded and averaged for statistical analysis.

### Assessment of motor coordination (Rotarod assay)

Motor coordination was evaluated using a rotarod device (Ugo Basile). Mice were trained over two consecutive days with three 1-minute sessions each day at a constant speed of 5 rpm. On the third day, performance was assessed on an accelerating rotarod, increasing from 5 to 40 rpm over 5 minutes, with a maximum trial duration of 400 seconds. Each mouse underwent three trials with a minimum 10-minute rest between trials. The latency to fall was recorded for each trial and the average of the three trials was used for analysis.

### In vitro muscle force measurement

In vitro muscle force measurements were performed on isolated extensor digitorum longus (EDL) using the 1200A Isolated Muscle System (Aurora Scientific). Muscles were maintained in an organ bath at 30°C containing oxygenated Ringer solution (137 mM NaCl, 24 mM NaHCO₃, 11 mM glucose, 5 mM KCl, 2 mM CaCl₂, 1 mM MgSO₄, and 1 mM NaH₂PO₄) continuously bubbled with 95% O₂/5% CO₂. After adjustment to the optimal muscle length (L₀), muscles were stimulated with 15 V electrical pulses to record peak twitch force (Pt).

Peak tetanic force (Po) was determined as the maximal force during a 500 ms stimulation at frequencies ranging from 10 to 250 Hz. Specific twitch and tetanic forces were calculated by normalizing force to the muscle cross-sectional area (CSA), using the formula: CSA (mm²) = muscle wet weight (mg) / [fiber length (L_f, mm) × 1.06 mg/mm³], where fiber length was estimated as L_f = L₀ × 0.44 for EDL as previously described.^81^

### Study design and statistical analysis

Mice were randomly assigned to experimental groups. Assessments of immunohistochemistry, muscle histology, muscle function, and behavioral assays were conducted by investigators blinded to group allocation. For statistical analysis, comparisons between two groups were performed using unpaired, two-tailed Student’s *t*-tests. For comparisons involving more than two groups, one-way ANOVA followed by either Bonferroni’s or Dunnett’s post hoc post-hoc test was applied. Normal distribution of variables was assumed for all analyses. Statistical tests were conducted using GraphPad Prism version 9.

## ACKNOWLEDGMENTS

We thank Filippo Oliveri and Lena Jörin for technical assistance; Dr. Alexander Ham and Dr. Marco Thürkauf for their support with AAV production; Dr. Alexander Schmidt and Dr. Thomas Bock for mass spectrometry analyses; and Dr. Dirk Grimm for providing the AAVMYO plasmids. This work was supported by the Cantons of Basel-Stadt and Basel-Landschaft, the Swiss Foundation for Research on Muscle Disease (FSRMM), Innosuisse (grant no. 34870.1 IP-LS), Santhera Pharmaceuticals, and the Muscular Dystrophy Association (grant no. https://doi.org/10.55762/pc.gr.157038).

## AUTHOR CONTRIBUTIONS

J.R.R. and M.A.R. conceived the project and designed the experiments. J.R.R. performed the majority of the experiments and analyzed the data. S.L. conducted histological quantifications and in vitro muscle force measurements. E.M. performed the RNA-seq data analysis. D.J.H. contributed to in vitro muscle force measurements. J.R.R. and M.A.R. wrote the manuscript and all authors provided feedback and approved the final version.

## DECLARATION OF INTERESTS

J.R.R. and M.A.R. are co-founders and shareholders of SEAL Therapeutics AG and are inventors on a patent application filed by the University of Basel related to this work.

