## Supplementary material for "Dual AAV gene therapy using laminin-linking proteins ameliorates muscle and nerve defects in LAMA2-related muscular dystrophy"

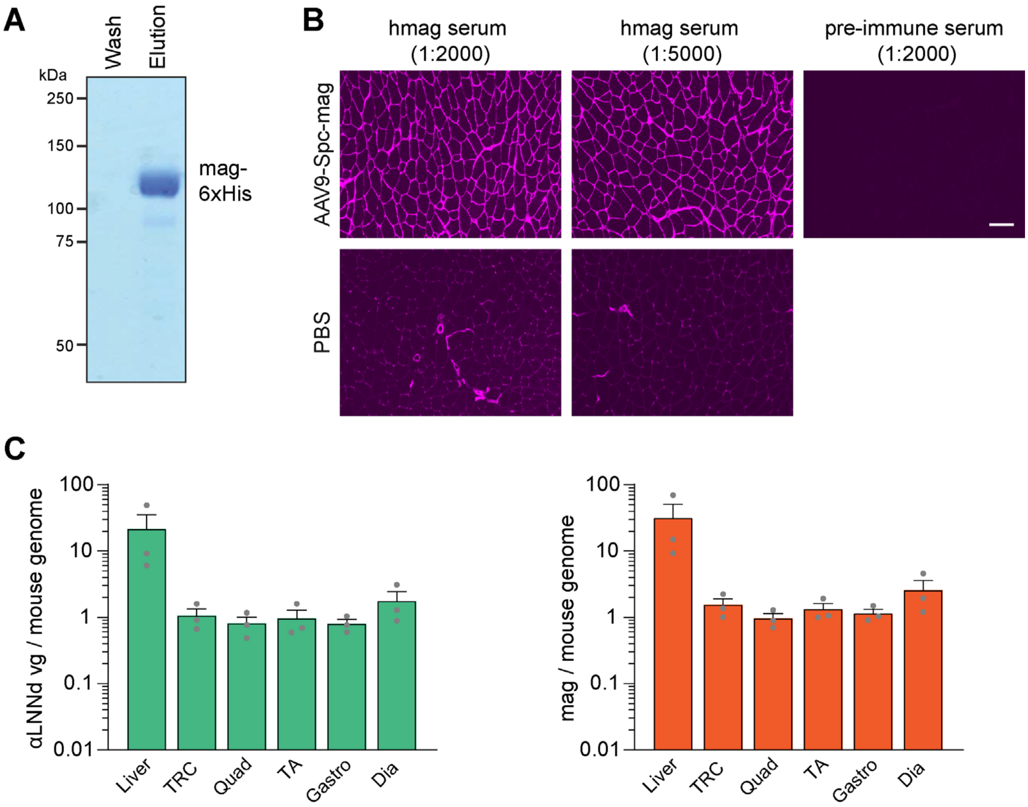


**Figure S1. Generation of anti-human mag antisera and assessment of AAV9-Spc-DL systemic transduction efficacy.**
(**A**) Coomassie Blue-stained SDS-PAGE of wash and eluate fractions from Ni-NTA columns loaded with supernatant from transiently transfected HEK293 cells expressing mag-6×His. (**B**) Immunofluorescence images of triceps muscle cross-sections from 8-week-old C57BL/6 mice injected intravenously at P1 with AAV9-Spc-*mag* ($1.5\times10^{14}$ vg/kg) or vehicle (PBS). Sections were stained using serum (various dilutions) from a rabbit immunized with *mag* or with pre-immune serum from the same rabbit. (**C**) Quantification of vector genomes (vg) normalized to mouse genomic DNA by qPCR from triceps muscles of 8-week-old C57BL/6 mice injected at P1 with AAV9-Spc-αLNNd and AAV9-Spc-mag ($1.5\times10^{14}$ vg/kg per construct). Data are presented as mean ± SEM. Scale bar: 100 μm. *n* = 3 mice per group.

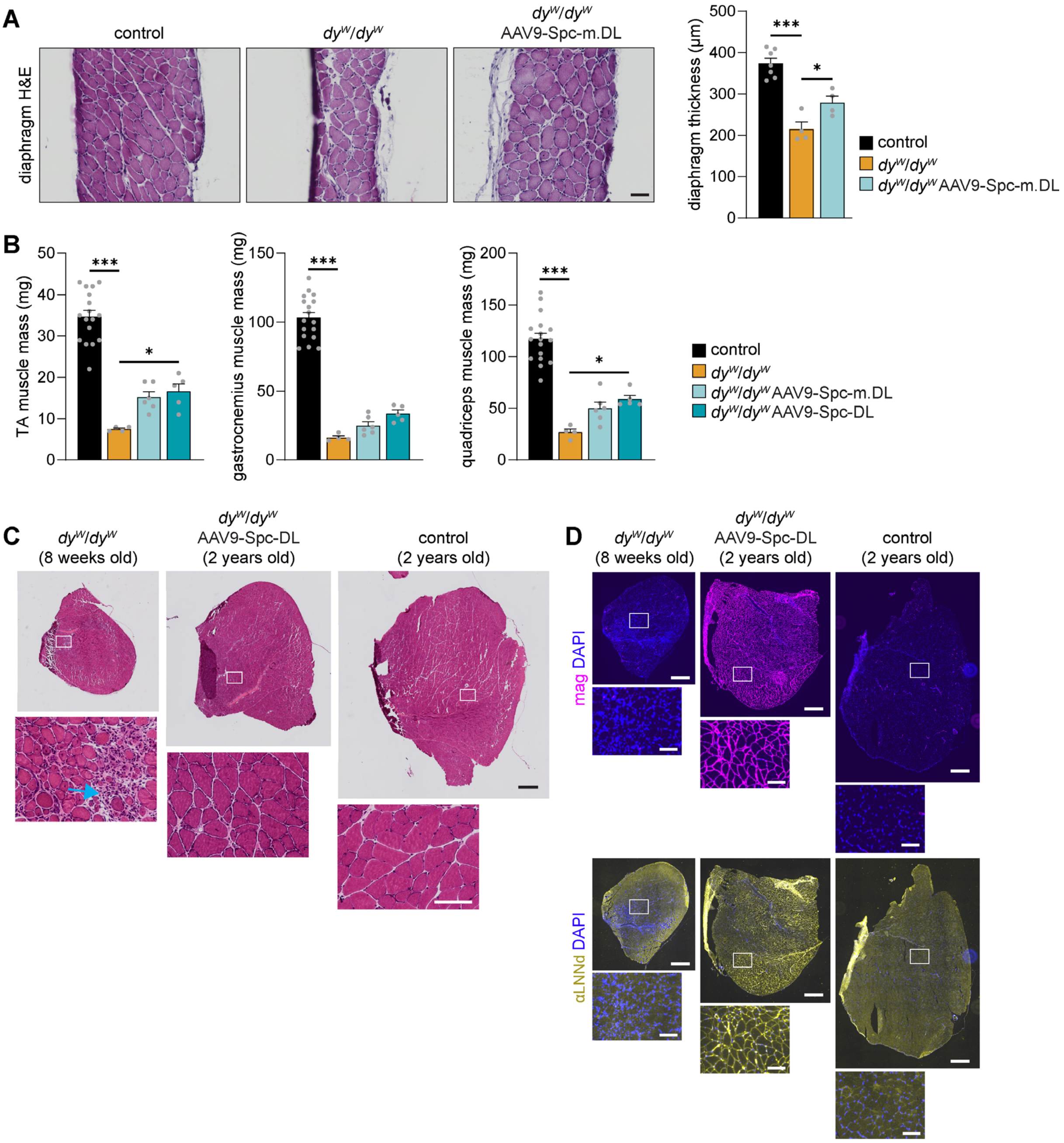


**Figure S2. Long-term expression and sustained therapeutic benefit of AAV-mediated, muscle-targeted expression of linker proteins in $dy^W/dy^W$ mice.**

(**A**-**D**) Histology, muscle mass and immunofluorescence of non-injected healthy control, $dy^W/dy^W$ mice, and $dy^W/dy^W$ mice injected intravenously at P1 with AAV9-Spc-m.DL or the corresponding human linker constructs (AAV9-Spc-DL) ($1.5\times10^{14}$ vg/kg per construct). (**A**) Representative H&E-stained cross-sections of the diaphragm from 8-week-old mice and quantification of diaphragm thickness. (**B**) Muscle mass in 8-week-old mice. (**C**-**D**) Representative images of triceps muscle cross-sections from an 8-week-old non-treated $dy^W/dy^W$ mouse, a 2-year-old treated $dy^W/dy^W$ and 2-year-old healthy control. (**C**) H&E staining. Notably, muscle from untreated mice $dy^W/dy^W$ showed mononuclear cell infiltrates (arrow), which were absent in treated $dy^W/dy^W$ mice. (**D**) Immunofluorescence images stained for αLNNd and mag. Data are presented as mean ± SEM. *$P < 0.05$, **$P < 0.01$, ***$P < 0.001$ by one-way ANOVA with Dunnett's post hoc test, comparing each group to untreated $dy^W/dy^W$ mice. Scale bars: 50 µm (A), 500 µm (C and D, top) and 100 µm (C and D, bottom). $n$ = 4-13 mice per group.

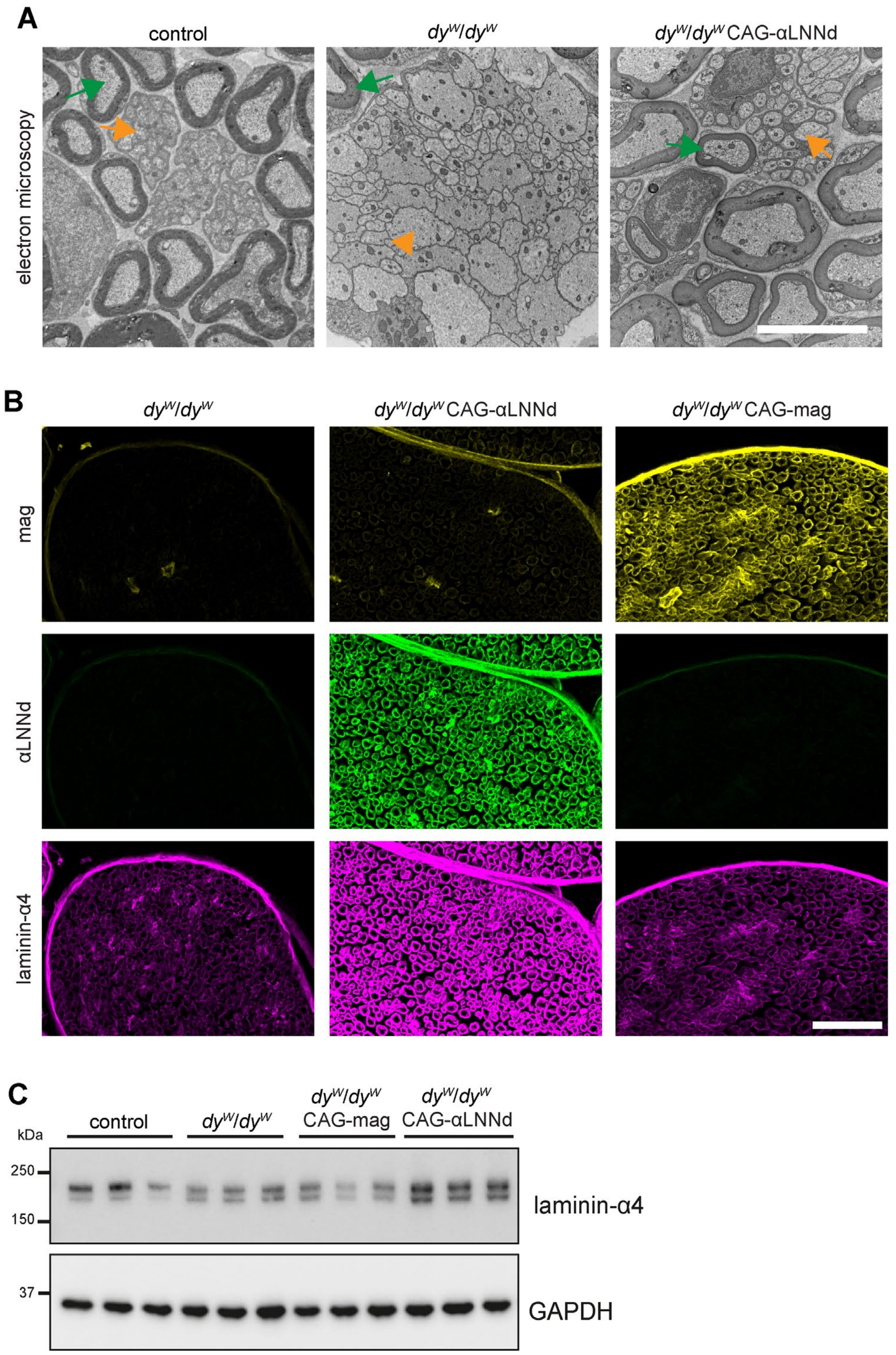


**Figure S3. Restoration of axonal sorting and increase in laminin-α4 in sciatic nerve following ubiquitous transgenic expression αLNNd.**

(**A-D**) Analysis of sciatic nerve pathology and laminin-α4 levels in 8-week-old healthy control mice, *dy*$^{W}$/*dy*$^{W}$ mice, or *dy*$^{W}$/*dy*$^{W}$ transgenic mice ubiquitously expressing mag or αLNNd under the control of the CAG promoter (CAG-mag or CAG-αLNNd). (**A**) Representative images of transmission electron microscopy of sciatic nerve from mice with the indicated genotypes. Note that in healthy controls, large-caliber motor axons are typically myelinated (green arrows), whereas small-diameter sensory axons are unmyelinated, ensheathed by non-myelinating Schwann cells, and clustered in Remak bundles (orange arrows). In contrast, *dy*$^{W}$/*dy*$^{W}$ mice display large regions of unsorted, non-myelinated axons, including large-caliber axons (orange arrowhead) not ensheathed by myelinating Schwann cells. In *dy*$^{W}$/*dy*$^{W}$ mice expressing CAG-αLNNd, axonal sorting appears similar to that observed in healthy controls with myelinated large axons (green arrow) and Remak bundels (orange arrow). (**B**) Immunofluorescence images of sciatic nerve cross-sections stained for mag, αLNNd and laminin-α4. Please note the strong increase in laminin-α4 in *dy*$^{W}$/*dy*$^{W}$ CAG-αLNNd mice compared to the other two genotypes. (**C**) Western blot analysis of laminin-α4 in sciatic nerve lysates. GAPDH was used as loading control. Scale bars: 5 µm (A), 50 µm (B).

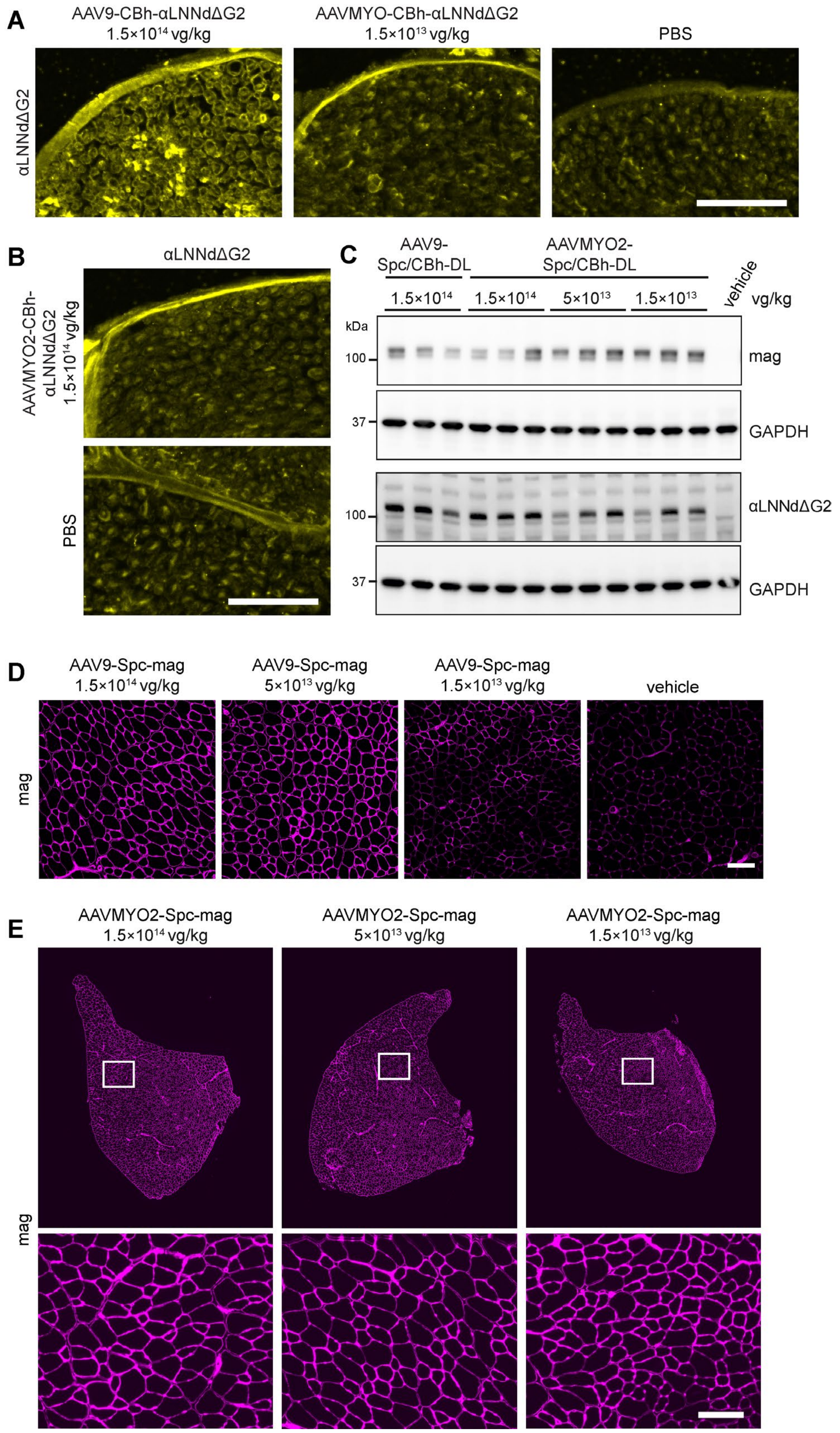


**Figure S4. Comparison of AAVMYO, AAVMYO2, and AAV9-mediated expression in sciatic nerve and skeletal muscle.**

(**A-D**) Analysis of linker protein amount in 8-week-old C57BL/6 mice injected intravenously at P1 with the indicated construct and doses. (**A-B**) Immunofluorescence images of sciatic nerve cross-sections stained for αLNNd. Notably, only AAV9-mediated transduction resulted in αLNNd-positive basement membranes within the endoneurium. Some background staining was also observed in axons of PBS-injected mice. (**C**) Western blot analysis of triceps muscle lysate to detect αLNNd or mag. GAPDH was used as loading control. (**D-E**) Immunofluorescence staining for mag of TA muscle cross-sections. Scale bars: 50 µm (A, B). 100 µm (D, E).

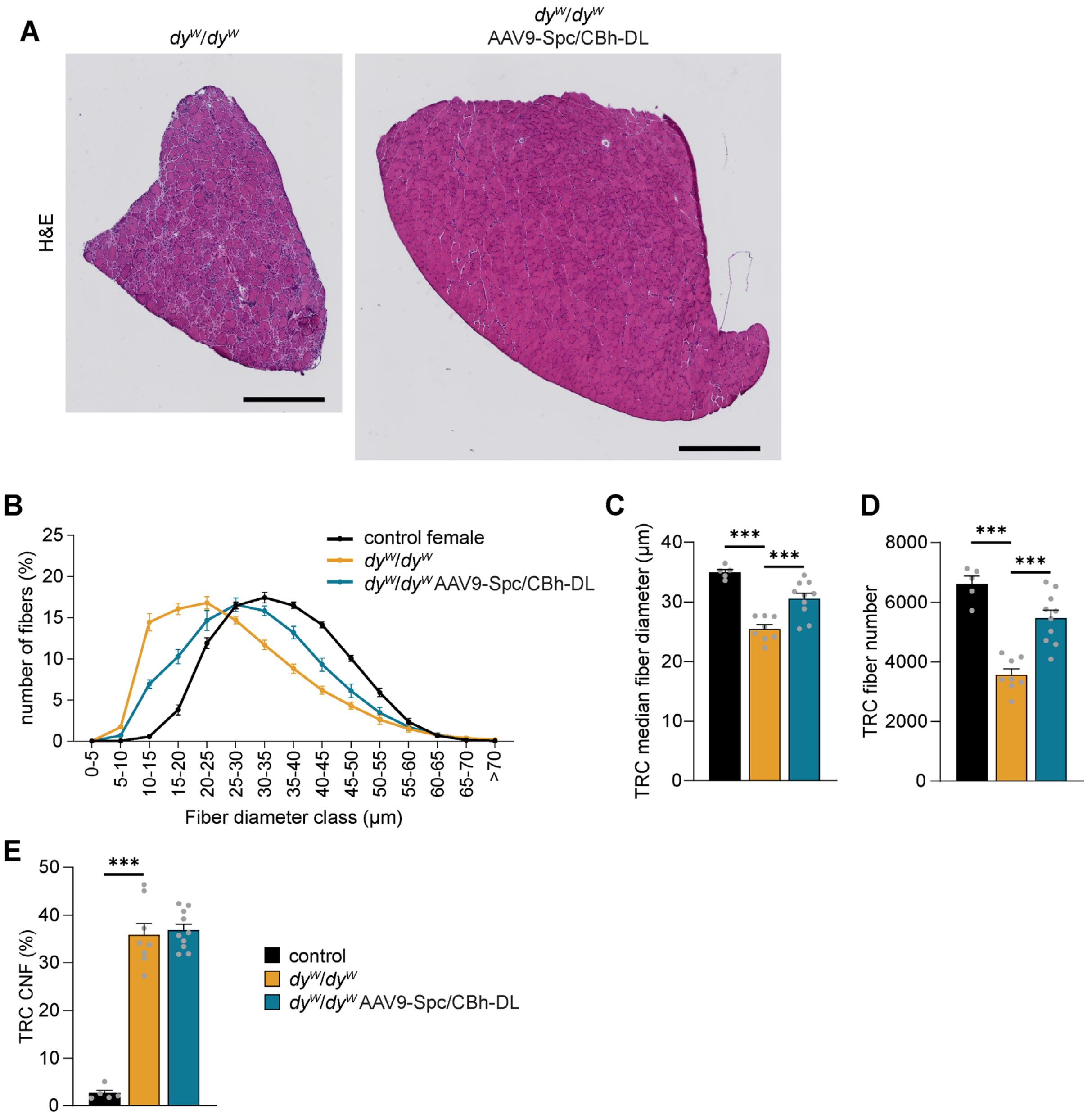


**Figure S5. AAV9-mediated improvement of histology in TA and triceps muscle in *dy^W/dy^W* AAV9-Spc/CBh-DL mice.**

(**A-E**) Histological analysis of skeletal muscle from 8-week-old wild-type, *dy^W/dy^W* and *dy^W/dy^W* mice injected intravenously at P1 with AAV9-Spc/CBh-DL ($1\times10^{14}$ vg/kg per construct). (**A**) Representative H&E-stained cross-sections of TA muscle. (**B-E**) Quantification of triceps muscle histology. (**B**) Distribution of muscle fiber diameters. (**C**) Median muscle fiber diameter. (**D**) Total number of muscle fibers. (**E**) Percentage of fibers with centralized nuclei. Data are presented as mean ± SEM. **P* < 0.05, ***P* < 0.01, ****P* < 0.001 by one-way ANOVA with Dunnett's post hoc test, comparing each group to untreated *dy^W/dy^W* mice. Scale bars: 500 *µ*m. *n* = 5-10 mice per group. Scale bar: 500 µm

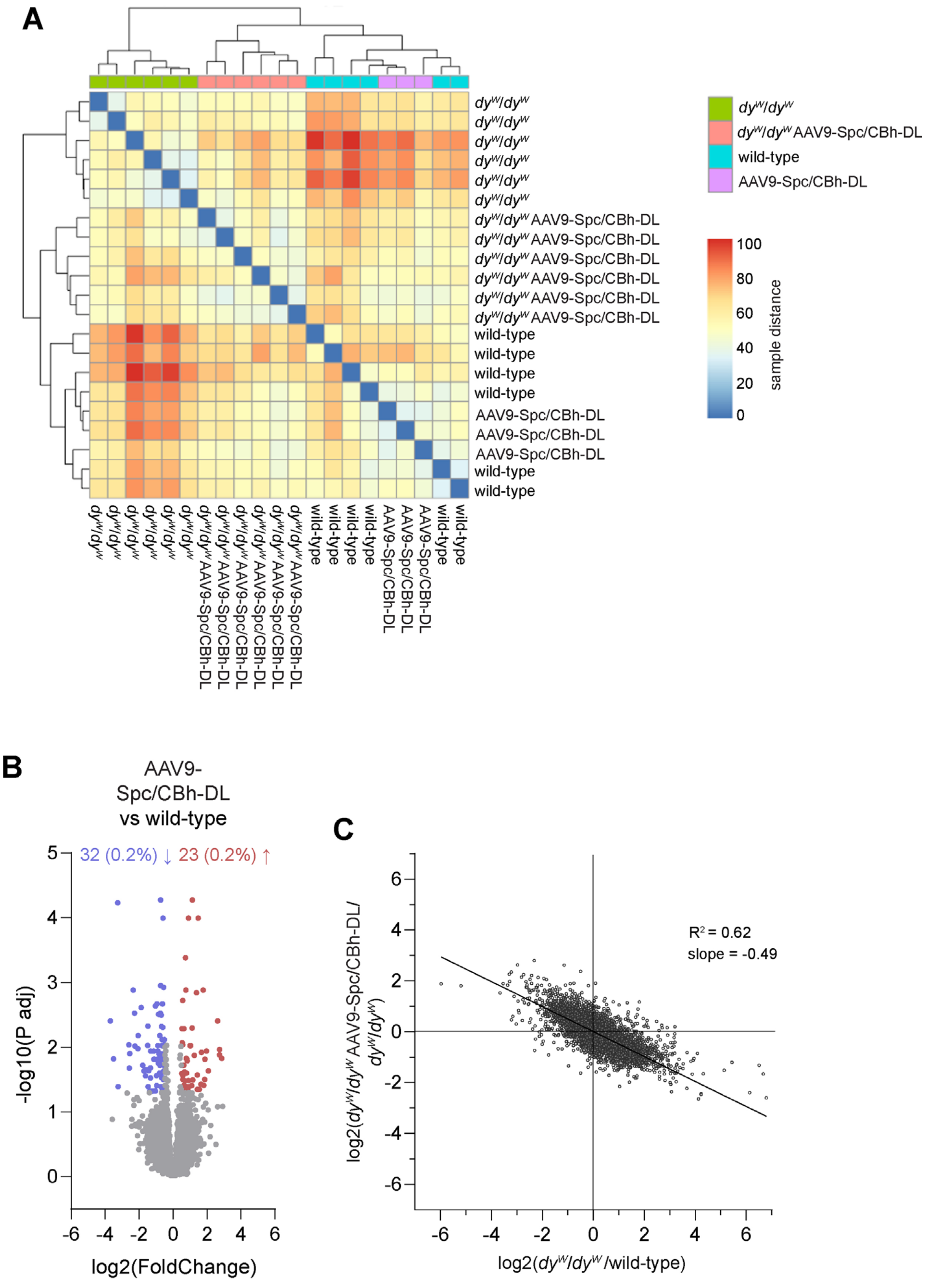


**Figure S6. RNAseq analysis of transcriptional changes by AAV9-mediated expression of mag and αLNNdΔG2 in *dy*$^{W}$/*dy*$^{W}$ mice and wild-type controls.**

(**A-E**) RNAseq analysis of TA muscle from 8-week-old mice: non-injected wild-type and *dy*$^{W}$/*dy*$^{W}$ mice, and wild-type and *dy*$^{W}$/*dy*$^{W}$ mice injected intravenously at P1 with AAV9-Spc/CBh-DL ($1\times10^{14}$ vg/kg per construct). (**A**) Sample correlation blot of RNAseq data. (**B**) Volcano plot showing the 14,905 transcripts detected in wild-type mice injected with AAV9-Spc/CBh-DL versus non-injected wild-type mice. Differentially expressed genes ($log_2$ fold change > ±0.5; adjusted $p < 0.05$) are shown in red (upregulated) or blue (downregulated). Numbers above indicate the total number and percentage of upregulated (red) or downregulated (blue) transcripts. (**C**) Pairwise comparison of gene expression changes: Fold change in *dy*$^{W}$/*dy*$^{W}$ versus wild-type mice plotted against fold change in *dy*$^{W}$/*dy*$^{W}$ AAV9-Spc/CBh-DL versus untreated *dy*$^{W}$/*dy*$^{W}$ mice. *n* = 3-6 mice per group.

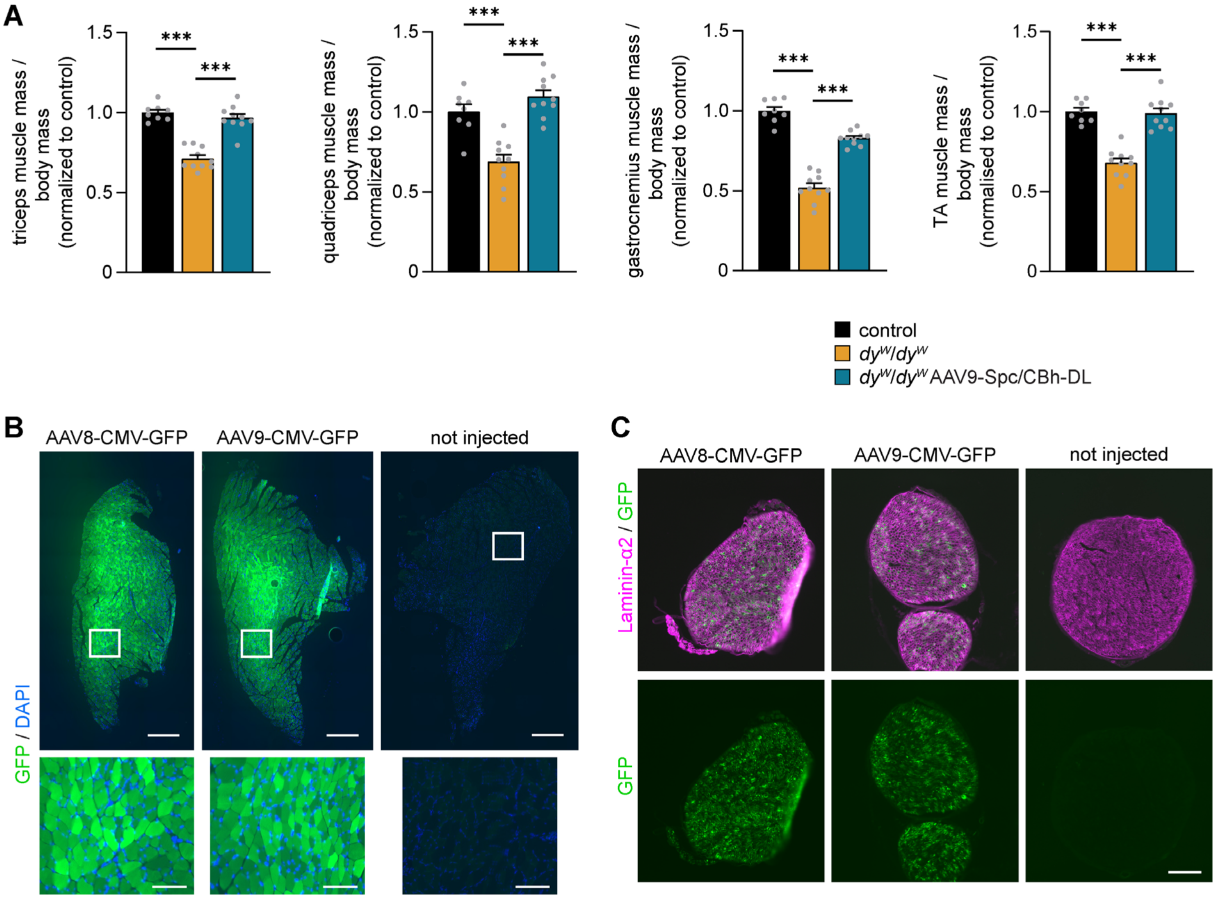


**Figure S7. Comparison of AAV9 and AAV8 transduction efficiency in muscle and peripheral nerve.** (**A**) Muscle mass normalized to body mass of 8-week-old wild-type, *dy$^{W}$/dy$^{W}$* or *dy$^{W}$/dy$^{W}$* mice intravenously injected at P1 with AAV9-Spc/CBh-DL ($1\times10^{14}$ vg/kg per construct). (**B-C**) C57BL/6 mice were injected intravenously at postnatal day 1 (P1) with CMV-GFP packaged into AAV9 or AAV8 ($1\times10^{14}$ vg/kg). Tissue was analyzed at 4 weeks of age. (**B**) Immunofluorescence images of TA muscle cross-sections stained for GFP. (**C**) Immunofluorescence images of sciatic nerve cross-sections stained for laminin-α2 and GFP. Scale bars: 500 µm (B, top) and 100 µm (B, bottom and C).

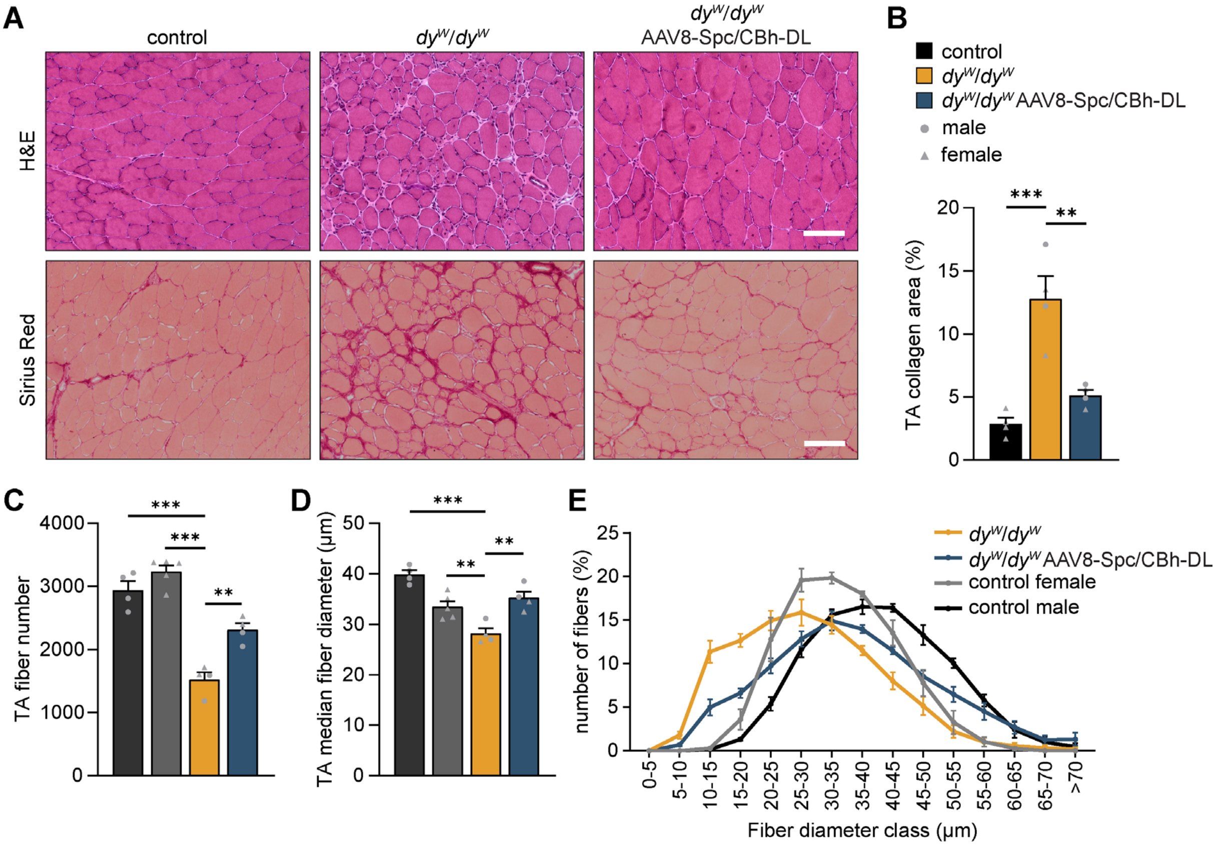


**Figure S8. Improvements in TA muscle histology by AAV8-mediated co-expression of mag in muscle and αLNNdΔG2 in muscle and nerve.**
(**A-E**) Analysis of TA muscle histology from 8-week-old wild-type, *dy$^{W}$/dy$^{W}$* or *dy$^{W}$/dy$^{W}$* mice injected at P1 with AAV8-Spc/CBh-DL ($1\times10^{14}$ vg/kg per construct). (**A**) Representative images of H&E- and Sirius Red-stained TA muscle cross-sections. (**B**) Quantification of muscle fibrosis based on Sirius Red-positive area in TA cross-sections. (**C**) Total muscle fiber number. (**D**) Median muscle fiber diameters. (**E**) Distribution of muscle fiber diameters. Data are presented as mean ± SEM. *$P < 0.05$, **$P < 0.01$, ***$P < 0.001$ by one-way ANOVA with Dunnett's post hoc test, comparing each group to untreated *dy$^{W}$/dy$^{W}$* mice. Scale bar: 100 µm. *n* = 4 mice per group.

**Video S1: Phenotype of 5-week-old *dy$^{W}$/dy$^{W}$* mice treated with AAV9-Spc-m.DL**

Representative video of a 5-week-old untreated *dy$^{W}$/dy$^{W}$* mouse and *dy$^{W}$/dy$^{W}$* mouse intravenously injected at P1 with AAV9-Spc-αLNNd and AAV9-Spc-mag (together referred to as AAV9-Spc-DL) at a dose of 1.5×$10^{14}$ vg/kg per construct.

**Video S2: Hindlimb paralysis in 8-month old *dy$^{W}$/dy$^{W}$* mouse treated with AAV9-Spc-m.DL**

Representative video of an 8-month-old *dy$^{W}$/dy$^{W}$* mouse intravenously injected at P1 with AAV9-Spc-DL (1.5×$10^{14}$ vg/kg per construct).

**Video S3: No signs of peripheral neuropathy in 8-month-old *dy$^{W}$/dy$^{W}$* mouse trangenically expressing αLNNd under a ubiquitous promoter**

Representative video of 8-month-old *dy$^{W}$/dy$^{W}$* CAG-αLNNd transgenic mouse showing normal gait and no evidence of peripheral neuropathy.

**Video S4: Peripheral neuropathy in 8-week-old *dy$^{W}$/dy$^{W}$* mouse trangenically expressing mag under a ubiquitous promoter**

Representative video of 8-week-old *dy$^{W}$/dy$^{W}$* CAG-mag transgenic mouse, displaying signs of peripheral neuropathy.

**Video S5: Close-to-normal phenotype in *dy$^{W}$/dy$^{W}$* mice treated with AAV9-Spc/CBh-DL.**

Representative video of 8-week-old wild-type, *dy$^{W}$/dy$^{W}$* and *dy$^{W}$/dy$^{W}$* mice injected intravenously at P1 with AAV9-Spc/CBh-DL (1×$10^{14}$ vg/kg per construct).

**Video S6: Improved neuromuscular phenotype by late treatment**

Representative movie of 12-week-old untreated *dy$^{W}$/dy$^{W}$* mice and *dy$^{W}$/dy$^{W}$* mice injected intravenously at 3 weeks of age with AAV8-Spc/CBh-DL (5×$10^{13}$ vg/kg per construct). Treated mice show improved motor performance.
